# Genetic effects on promoter usage are highly context-specific and contribute to complex traits

**DOI:** 10.1101/319806

**Authors:** Kaur Alasoo, Julia Rodrigues, John Danesh, Daniel F. Freitag, Dirk S. Paul, Daniel J. Gaffney

## Abstract

Genetic variants regulating RNA splicing and transcript usage have been implicated in both common and rare diseases. Although transcript usage quantitative trait loci (tuQTLs) have now been mapped in multiple cell types and conditions, the molecular mechanisms through which these variants exert their effect have remained elusive. Specifically, changes in transcript usage could arise from promoter choice, alternative splicing or 3′ end choice, but current tuQTL studies have not been able to distinguish between them. Here, we performed comprehensive analysis of RNA-seq data from human macrophages exposed to a range of inflammatory stimuli (IFNγ, *Salmonella*, IFNγ + *Salmonella*) and a metabolic stimulus (acetylated LDL), obtained from up to 84 individuals. In addition to conventional gene-level and transcript-level analyses, we also developed an analytical approach to directly quantify promoter, internal exon and 3′ end usage. We found that although naive transcript-level analysis often links single genetic variants to multiple coupled changes on the transcriptome, this appears to be an artefact of incomplete transcript annotations. Most of this coupling disappears when promoters, splicing and 3′ end usage are quantified directly. Furthermore, promoter, splicing and 3′ end QTLs are each enriched in distinct genomic features, suggesting that they are predominantly controlled by independent regulatory mechanisms. We also find that promoter usage QTLs are 50% more likely to be context-specific than canonical splicing QTLs and constitute 25% of the transcript-level colocalisations with complex traits. Thus, promoter usage might be a previously underappreciated molecular mechanism mediating complex trait associations in a context-specific manner.

## Introduction

Genome-wide association studies (GWAS) have discovered thousands of genetic loci associated with complex traits and diseases. However, identifying candidate causal genes and molecular mechanisms at these loci remains challenging. Complex trait-associated variants are enriched in regulatory elements and are therefore thought to act via regulation of gene expression levels, often in a cell type-and context-specific manner [1–3]. However, such variants are equally enriched among splicing quantitative trait loci (QTLs) [4,5] and incorporating splicing QTLs in a transcriptome-wide association study increased the number of disease-associated genes by 2-fold [6]. In addition to splicing, genetic variants can also alter transcript sequence by regulating promoter and 3′ end usage, which we refer to collectively hereafter as transcript usage QTLs (tuQTLs). Alternative transcript start and end sites underlie most transcript differences between tissues [7,8], they are dynamically regulated in response to cellular stimuli [9,10] and they are also frequently dysregulated in cancer [11,12]. Moreover, experimental procedures designed to capture either 5′ or 3′ ends of transcripts have identified disease-relevant genetic variants that regulate promoter or 3′ end usage [13,14]. However, well-powered RNA-seq-based tuQTL studies performed across cell types [5,6,15–17] and conditions [18–20] have thus far not distinguished between promoter usage, splicing and 3′ end usage. Thus, how these distinct transcriptional mechanisms contribute to complex traits and how context-specific these genetic effects are is currently unclear.

In addition to splicing analysis, RNA-seq data can also be used to quantify promoter and 3′ end usage. The simplest approach would be to first quantify the expression of all annotated transcripts using one of the many quantification algorithms (benchmarked in [21]). Linear regression can then be used to identify genetic variants that are associated with the usage of each transcript of a gene [6,16]. Comparing the associated transcripts to each other can reveal which transcriptional changes take place (Fig. 1a). A key assumption here is that all expressed transcripts are also part of the annotation catalog. If some of the expressed transcripts are missing, then reads originating from the missing transcripts might be erroneously assigned to other transcripts that are not be expressed at all (Fig. 1b) [22]. This can lead to individual genetic changes being spuriously associated with multiple transcriptional changes. For example, a genetic variant regulating promoter usage might also appear to be associated with the inclusion of an internal exon (Fig. 1b), although there are no reads originating from that exon. Importantly, this is not just a theoretical concern, because 25-35% of the exon-exon junctions observed in RNA-seq data are not present in transcript databases [16], and up to 60% of the transcripts annotated by Ensembl [23] are truncated at the 5′ or 3′ end (Figures S1 and S2).

**Fig. 1.**
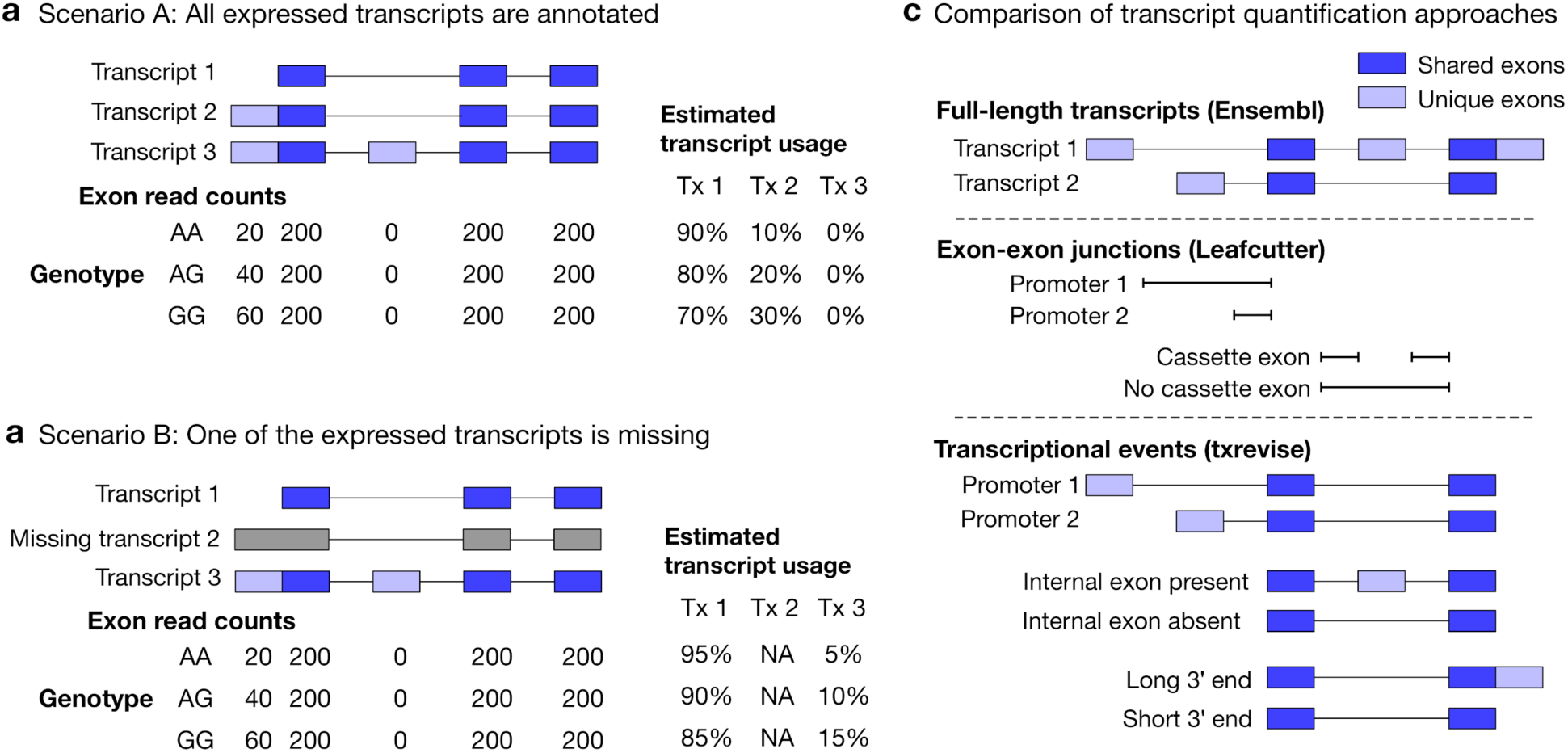
Challenges of quantifying transcript usage from RNA-seq data. Transcript quantification seeks to estimate the most likely configuration of *known* transcripts that best explains observed read counts supporting the inclusion of each exon. (**a**) In scenario A, each copy of the G allele increases the usage of transcript 2 by 10%. Since both expressed transcripts (transcript 1 and transcript 2) are annotated, we successfully detect the change and conclude that the G allele increases the expression of the proximal promoter of the gene. (**b**) In scenario B, each copy of the G allele still increases the usage of transcript 2 by 10%. However, since transcript 2 is missing from the annotations, reads originating from transcript 2 are now falsely assigned to transcript 3. Since transcript 3 also contains alternative second exon, we now falsely conclude that in addition to promoter usage, the G allele is also associated with increased inclusion of exon 2, even though there are no reads mapping to exon 2. Furthermore, the magnitude of the genetic effect is underestimated, because the reads assigned to transcript 3 are assumed to be evenly distributed across the promoter and the alternative exon. (**c**) Top panel: Two hypothetical transcripts that differ from each other at the promoter, at an internal exon and at the 3′ end. Middle panel: Leafcutter uses reads mapping to exon-exon junctions to identify alternatively excised introns. Bottom panel: txrevise uses the exons shared between transcripts (dark blue) as a scaffold to construct three independent transcriptional events from the two original transcripts.

The overcome the issue of missing transcript annotations, recent tuQTL studies have focussed on quantifying transcription at the level of individual exons [15,24,25], introns [25] or exon-exon junctions (Fig. 1c) [6,16,25]. While these approaches often discover complementary genetic associations [16,25], they do not explicitly reveal the transcriptional mechanism (promoter usage, alternative splicing or 3′ end usage) underlying the genetic associations. The most successful approach to differentiate between distinct transcriptional mechanisms has been ‘event-level’ analysis where reference transcripts are split into independent events (e.g. promoters, splicing events and 3′ end) whose expressions is then quantified using standard transcript quantification methods (Fig. 1c). This approach was pioneered by MISO [26] and was recently used to identify promoter usage QTLs in the GEUVADIS dataset [9]. Despite its success, MISO covers only a subset of promoter events (alternative first exons) and its event annotations have not been updated since it was first published. Thus, there is a need for a method that is able to detect comprehensive set of promoter, splicing and 3′ end usage QTLs in an uniform manner.

In this study, we re-analysed RNA-seq data from human macrophages exposed to three inflammatory stimuli (18 hours IFNγ stimulation, 5 hours *Salmonella* infection and IFNγ stimulation followed by *Salmonella* infection). We also collected a new dataset of macrophages stimulated with acetylated LDL (acLDL) for 24 hours. We mapped genetic associations at the level of total gene expression, full-length transcript usage and exon-exon junction usage in each experimental condition. In addition to existing quantification methods, we also developed a complementary approach (txrevise) that stratifies reference transcript annotations into independent promoter, splicing and 3′ end events. Using txrevise, we found that promoter and 3′ end usage QTLs constituted 55% of detected tuQTLs, exhibited genetic architectures that were distinct from canonical expression or splicing QTLs and often colocalised with complex trait associations. Promoter usage QTLs were also 50% more likely to be context-specific than canonical splicing QTLs. Thus, context-specific regulation of promoter usage might be a previously underappreciated molecular mechanism underlying complex trait associations.

## Results

### Quantifying transcript usage in stimulated macrophages

We analysed RNA-seq data from human induced pluripotent stem cell (iPSC)-derived macrophages exposed to three inflammatory stimuli (18 hours IFNγ stimulation, 5 hours *Salmonella* infection, and IFNγ stimulation followed by *Salmonella* infection) and one metabolic stimulus (24 hours acLDL stimulation). While the gene expression analysis of the IFNγ + *Salmonella* dataset from 84 individuals has previously been described [3], the acLDL data from 70 individuals was newly generated for the current study. The acLDL dataset allowed us to assess how our results generalise to weaker non-inflammatory stimuli. Both datasets included independent unstimulated control samples (denoted as ‘naive’ and ‘Ctrl’). In each condition, we quantified gene expression and transcript usage using the following established quantification approaches: (i) gene-level read count quantified with featureCounts [27], (ii) full-length transcript usage quantified with Salmon [28] (Fig. 1c), and (iii) exon-exon junction usage quantified with Leafcutter [6] (Fig. 1c).

Inspired by event level analysis proposed by MISO [9,26], we also developed a complementary approach (txrevise) to stratify reference transcript annotations into independent promoter, splicing and 3′ end events. To achieve this, txrevise identifies constitutive exons shared between all transcripts of a gene and uses those to assign non-constitutive exons to promoter, internal exon or 3′ end events (Fig. 1c). Since up to 60% of the transcripts annotated by Ensembl [23] are truncated at the 5′ or 3′ end (Figure S1), txrevise extends truncated transcripts by copying over exons from the longest transcript of the gene (Figure S2). This step eliminates a large number of implausible alternative promoters and 3′ ends that lack experimental evidence. To make the approach suitable for genes with non-overlapping transcripts, we also select a subset of transcripts that share the largest number of exons between them (Figure S3). Finally, to ensure that the new alternative promoter and 3′ end events do not capture splicing changes, txrevise masks alternative exons that are not the first or last exons (Figure S4). The R package as well as custom transcriptional events constructed by txrevise are available from GitHub (https://github.com/kauralasoo/txrevise).

### Genetic effects on transcript usage

Depending on the experimental condition and quantification method, we detected between 1500 and 3500 QTLs at the 10% false discovery rate (FDR) (Fig. 2a). Leafcutter consistently detected the lowest number of QTLs per condition, while txrevise detected approximately 30% more associations than other methods (Fig. 2a), 55% of which affected promoter or 3′ end usage instead of internal exons (Figure S5). However, this increase in QTLs can be partially explained by the fact that txrevise detected multiple associations for ∼24% of the genes while the full-length tuQTL analysis was limited to single lead association per gene (Figure S6). Some of these additional QTLs are likely to represent independent causal variants, such as the three independent tuQTLs detected for the *IRF5* gene (Figure S7). Alternatively, as discussed below, additional associations could also be caused by transcriptional coupling where promoter or 3′ end choice directly influences the splicing of an internal exons or *vice versa [29]*.

**Fig. 2.**
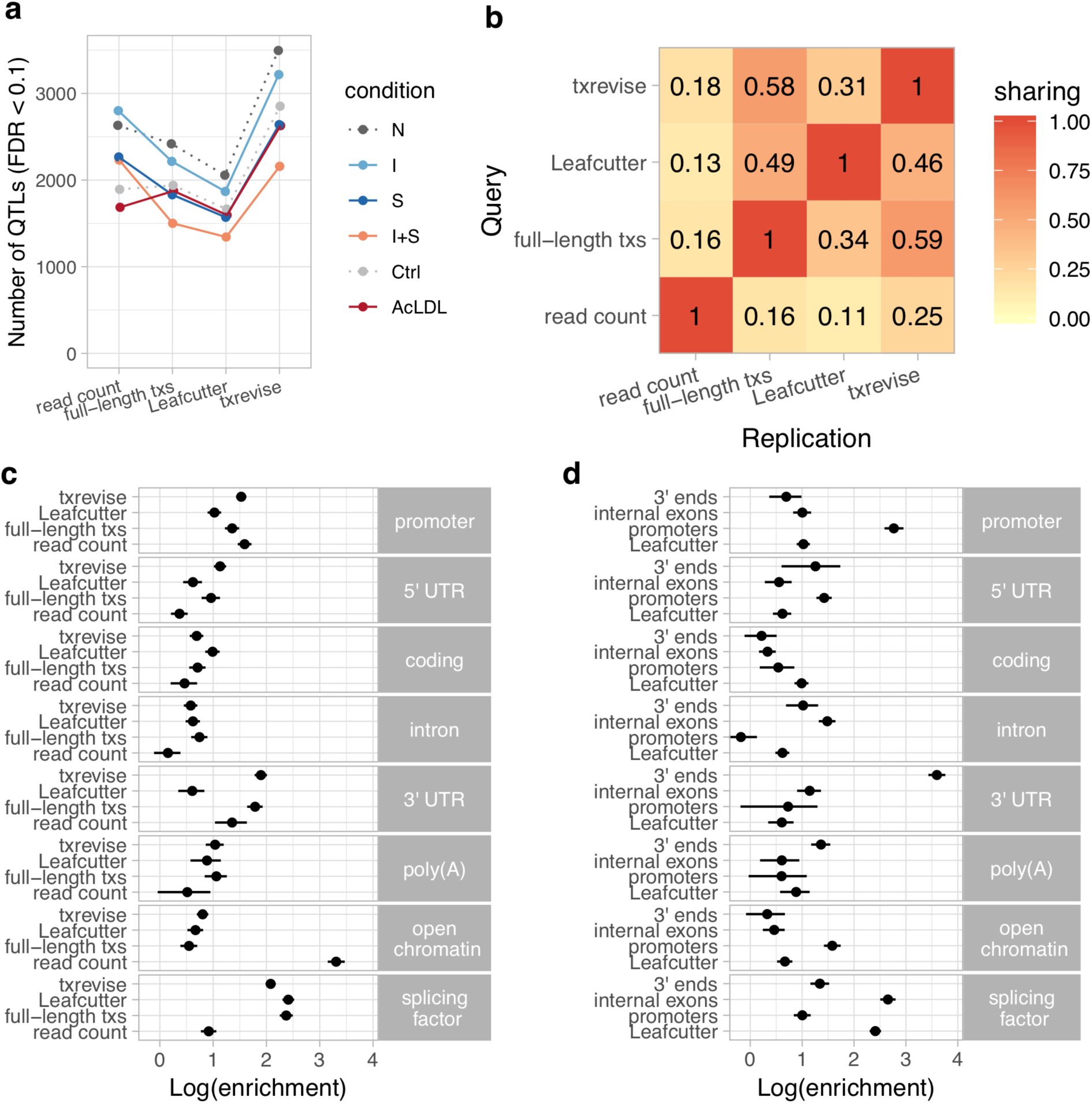
Diversity of QTLs detected by different quantification methods. In panels a-c, all txrevise QTLs from promoters, internal exons and 3′ ends have been pooled to facilitate comparison with eQTLs as well as Leafcutter and full-length transcript usage QTLs. (**a**) Number of QTLs detected by read count, full-length transcript usage, Leafcutter and txrevise methods in each condition (N, naive; I, IFNγ; S, *Salmonella*, I+S, IFNγ + *Salmonella*). (**b**) Sharing of QTLs detected by four quantification methods. The numbers on the heatmap show the fraction of QTLs detect by one method that were replicated by the three other methods (R^2^ > 0.8 between lead variants). Only QTLs with FDR < 0.01 were included in the analysis. (**c**) Enrichment of genomic annotations at QTLs detected by the four quantification methods. (**d**) Comparison of Leafcutter tuQTLs to promoter, internal exon and 3′ end usage QTLs detected by txevise. Genomic annotations used for enrichment analysis: *promoter* - promoter flanking regions (−2000 bp to +200 bp); *5′ UTR, coding, intron, 3′ UTR* - corresponding regions extracted from Ensembl transcripts; *poly(A)* - experimentally determined polyadenylation sites (+/- 25 bp) [31]; *open chromatin* - open chromatin regions from macrophages [3]; *splicing factor* - experimentally determined binding sites of splicing factors detected by eCLIP [32]. The points on panels c and d show the natural logarithm of enrichment for each annotation and the lines represent the 95% confidence intervals from fgwas [33].

Different quantification methods may be biased towards discovering events with specific genomic properties, which is not captured by the number of QTLs detected. To address this, we quantified how often were the lead QTL variants (FDR < 0.01) from different methods in high linkage disequilibrium (LD) (R^2^ > 0.8) with each other (see Methods). Consistent with previous reports that tuQTLs are largely independent from eQTLs [5], we found that only 11-24% of the lead variants detected at the read count level replicated at the transcript level (R^2^ > 0.8, irrespective of the replication p-value), independent of which quantification method was used (Fig. 2b). In contrast, ∼50% of the Leafcutter QTLs were also detected by txrevise or full-length transcript usage approaches. Similarly, the tuQTLs detected by txrevise and full-length transcript usage quantification were in high LD more than 60% of the time (Fig. 2b). Finally, we found that while 44% of the txrevise internal exon QTLs were in high LD with Leafcutter QTLs, this decreased to ∼20% for promoter and 3′ end QTLs (Figure S5), suggesting that Leafcutter is less suited to capture those events. Thus, different quantification approaches appear to capture complementary sets of genetic associations.

### Transcriptional coupling between promoter usage, splicing and 3′ end usage

Although multiple instances of transcriptional coupling have been previously observed (reviewed in [29]), its genome-wide prevalence is still poorly understood. Proximal genetic variants predominantly regulate transcript usage in *cis*. Thus, a single causal variant associated with multiple transcriptional events could provide evidence for transcriptional coupling in the absence of full-length mRNA sequencing data [30]. To test this, we first focussed on the tuQTLs detected by full-length Ensembl transcripts. We compared the transcript with the smallest tuQTL p-value against the most highly expressed transcript (see Methods). This revealed that 97% of the tuQTLs were seemingly associated with at least two transcriptional changes (e.g. alternative promoter and internal exon) and >50% were simultaneously associated with all three transcriptional changes (different promoter, internal exons and 3′ end) (Figure S8). In contrast, a recent full-length mRNA sequencing experiment estimated coupling to occur between approximately 10% of the events [30], suggesting that our naive estimate is strongly inflated by the use of incomplete transcript annotations (Fig. 1a, b).

To estimate the extent to which incomplete transcript annotations can cause false signals of transcriptional coupling, we turned to txrevise and Leafcutter analyses that directly quantified the usage of multiple independent transcriptional events per gene. Both approaches detected multiple significant tuQTLs for only a minority of the genes (txrevise −24%; Leafcutter - 10%) (Figure S6). Furthermore, among the genes with multiple tuQTLs, 50% of the Leafcutter and 30% of the txrevise associations had independent lead variants (R^2^ < 0.2), confirming that true transcriptional coupling seems to be rare (Figure S6). Moreover, some of the apparent coupling events in the txrevise analysis could be explained by technical biases such as large eQTL effects (Figure S9) or positional biases in the RNA-seq data (Figure S10). Our results suggest that, although most genetic variants modulate individual transcriptional events, high numbers of spurious associations between genotype and multiple transcriptional changes are detected when expression is quantified at the level of full-length transcripts.

### Genomic properties of transcript usage QTLs

To characterise the genetic associations detected by different quantification methods, we compared the relative enrichments of the identified QTLs across multiple genomic annotations. We constructed genomic tracks for eight annotations: open chromatin measured by ATAC-seq [3], promoter flanking regions (−2000 bp to +200 bp), 5′ UTRs, coding sequence (CDS), introns, 3′ UTRs, polyadenylation sites [31], and eCLIP binding sites for RNA binding proteins involved in splicing regulation (splicing factors) [32]. We then used the hierarchical model implemented in fgwas [33] to estimate the enrichment of each genomic annotation among the QTLs detected by each quantification method. Consistent with the limited overlap between eQTLs and tuQTLs (Fig. 2b), we found that eQTLs were strongly enriched in sites of open chromatin (Fig. 2c; log enrichment of 3.31, 95% CI [3.15, 3.47]), whereas all transcript-level QTLs were enriched at the binding sites of splicing factors detected by eCLIP (Fig. 2c, mean log enrichment of 2.29). Importantly, when all txrevise tuQTLs were pooled, the enrichment patterns were broadly similar to tuQTLs detected by full-length Ensembl transcripts (Fig. 2c). This suggests that txrevise events and full-length transcripts capture similar genetic associations but txrevise facilitates more accurate identification of the underlying transcriptional event (i.e. promoter, internal exon or 3′ end usage) (Fig. 2b). Finally, compared to Leafcutter, full-length transcript usage and txrevise QTLs were both more strongly enriched at 3′ UTRs (Fig 2c, mean log enrichment of 1.85), suggesting that they capture changes in 3′ UTR length that do not manifest at the level of junction reads and are thus missed by Leafcutter.

If the coupling between promoter usage, splicing and 3′ end choice is as rare as suggested by txrevise, then this should be reflected by the genomic features that the associated variants are enriched in. Conversely, pervasive coupling between distinct transcriptional mechanisms would predict that the associated variants would share most of their genomic properties. To assess this, we repeated the fgwas analysis on the promoter, internal exon and 3′ end QTLs detected by txrevise as well as Leafcutter splicing QTLs. We found that Leafcutter and internal exon QTLs showed broadly similar enrichment patterns, with a strong enrichment at the binding sites of splicing factors (Fig. 2d, mean log enrichment of 2.53). In contrast, promoter and 3′ end usage QTLs were specifically enriched at promoters (Fig. 2d; log enrichment of 2.76, 95% CI [2.59, 2.95]) and 3′ UTRs (Fig 2d; log enrichment of 3.60, 95% CI [3.43, 3.76]), respectively (Fig. 2d), and showed only a modest enrichment at the binding sites of splicing factors (Fig. 2d; mean log enrichment of 1.17). Compared to other events, promoter usage QTLs were relatively more enriched in open chromatin regions (log enrichment of 1.58, 95% CI [1.42, 1.74]) and were also 65% more likely to overlap chromatin accessibility QTLs in macrophages [3] (R^2^ > 0.9) (Fisher’s exact test p-value = 3.87×10^-5^). Thus, promoter usage, splicing and 3′ end usage appear to be regulated by largely independent sets of genetic variants enriched in distinct genomic regions.

### Colocalisation with complex trait associations

To assess the relevance of different QTLs for interpreting complex trait associations, we performed statistical colocalisation analysis with GWAS summary statistics for 33 immune-mediated and metabolic traits and diseases (see Methods). We found that 47 of 138 colocalised QTLs influenced total gene expression level (Fig. 3a) (PP3+PP4>0.8, PP4/PP3>9; PP3, posterior probability of a model with two distinct causal variants; PP4, posterior probability of a model with one common causal variant). In contrast, the remaining 91 colocalised QTLs were associated with at least one of the transcript-level phenotypes (full-length transcript usage, txrevise or Leafcutter) but not with total gene expression (Fig. 3a). Similarly, 44 of 91 transcript-level colocalisations were detected only by a single transcript quantification approach (Fig. 3a). An important caveat of this analysis is that it does not directly test if the colocalisations are specific to one quantification method or simply missed by others because of limited power. Thus, our estimates of method-specificity are likely to be inflated.

**Fig. 3.**
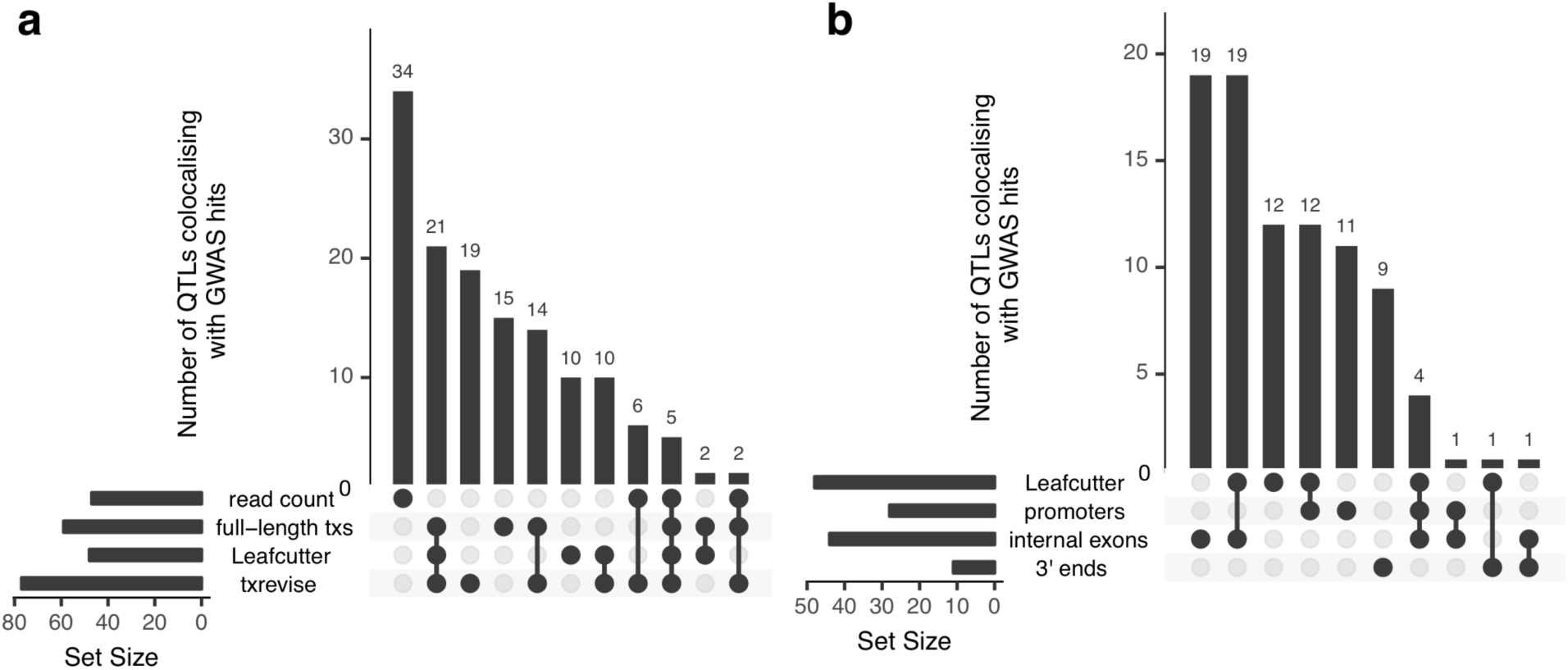
Colocalisation between GWAS hits for 33 complex traits and QTLs. The UpSetR plot shows the total number of colocalised trait-gene pairs detected by different quantification methods (horizontal bars) and overlap between different methods (vertical bars). (**a**) Sharing of colocalisations between four quantification methods. (**b**) Sharing of colocalisations between Leafcutter and three independent txrevise event types (promoters, internal exons, 3′ ends).

Finally, to quantify the relative contribution of promoter usage, splicing and 3′ end usage to complex traits, we stratified the txrevise colocalisations by transcriptional event type. We found that 44 of 77 colocalised QTLs influenced internal exons and the rest regulated promoters and 3′ ends (Fig. 3b). We were able to replicate known associations between splicing of exon 2 in *CD33* and Alzheimer’s disease (Figure S11) [34] and splicing of exon 13 in *HMGCR* and LDL cholesterol (Figure S12) [35]. Importantly, while half of the promoter and internal exon colocalisations were also detected by Leafcutter, only 1/11 3′ end events were captured by Leafcutter, probably because these are less likely to manifest at the level of junction reads (Fig. 3b).

### Condition-specificity of expression and transcript usage QTLs

Next, we explored how the genetic effects of eQTLs and tuQTLs varied in response to stimuli. To define response QTLs, we started with QTLs detected (FDR < 10%) in each of the four simulated conditions (I, S, I+S and acLDL) and used an interaction test to identify cases where the QTL effect size was significantly different between the simulated and corresponding naive condition (FDR < 10%). To exclude small but significant differences in effect size, we used a linear mixed model to identify QTLs where the interaction term explained more than 50% of the total genetic variance in the data (see Methods). Although the fraction of QTLs that were response QTLs varied greatly between conditions (Fig. 4a) and correlated with the number of differentially expressed genes (Figure S13) as previously reported [1], we found that the fraction of response tuQTLs was relatively consistent between the four quantification methods (Fig. 4a). While previous reports have highlighted that eQTLs are more condition-specific than tuQTLs [18], we found no clear pattern in our data with stronger stimuli (S and I+S) showing larger fraction of condition-specific eQTLs, and weaker stimuli (I, acLDL) showing smaller fraction of response eQTLs (Fig. 4a) compared to tuQTLs. However, when we focussed on the transcriptional events detected by txrevise, we found that promoter usage QTLs were 50% more likely to be response QTLs than tuQTLs regulating either internal exons or 3′ ends (Fig. 4b) (Fisher’s exact test combined p-value = 2.79×10^-6^).

**Fig. 4.**
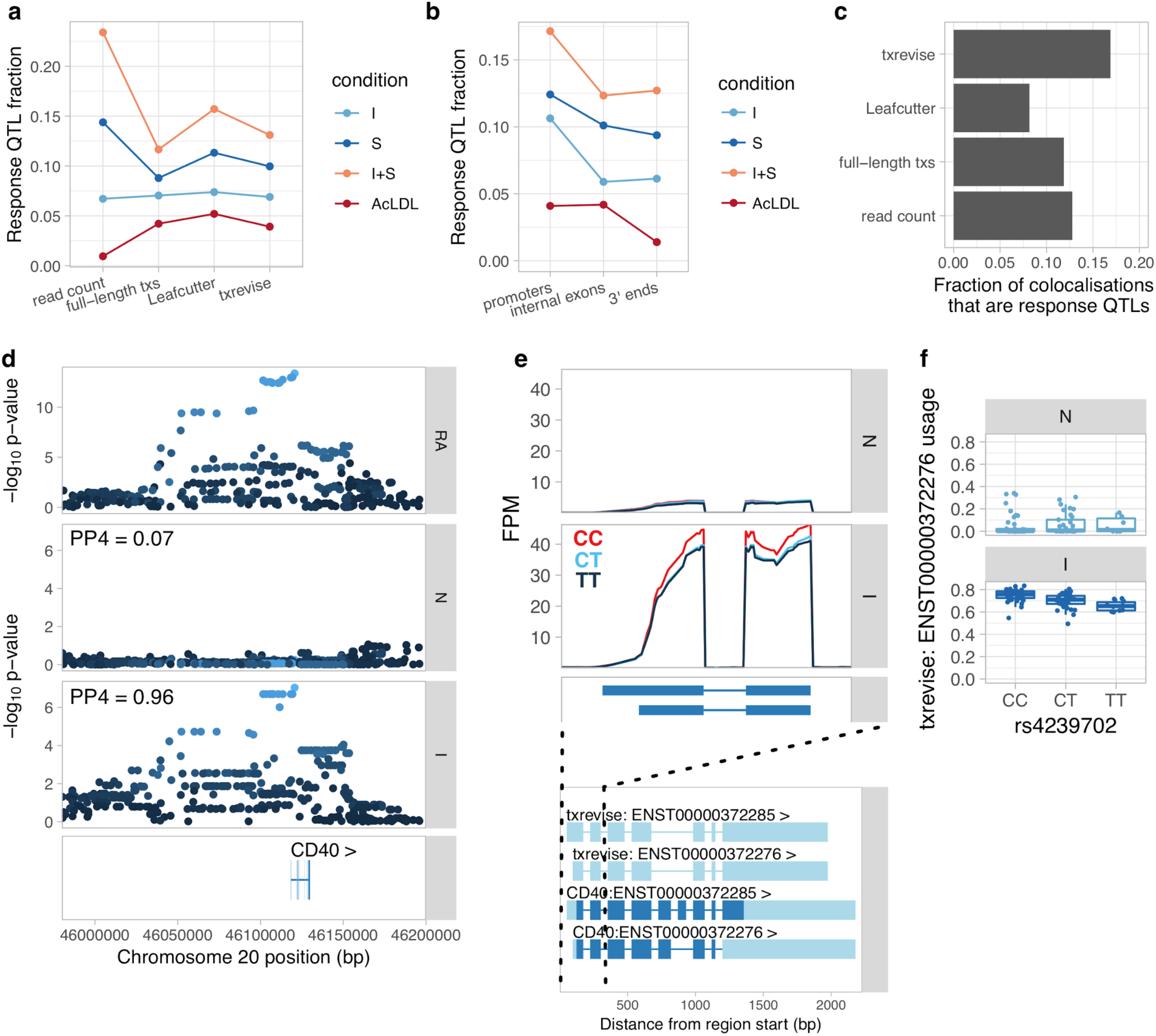
Condition-specificity of eQTLs and tuQTLs. (**a**) Fraction of all QTLs detected in each simulated condition that are response QTLs (FDR < 10% and more than 50% of the genetic variance explained by the interaction term). (**b**) Fraction of txrevise tuQTLs classified as response QTLs, stratified by the part of the gene that they influence (promoters, internal exons or 3′ ends). (**c**) Fraction of GWAS colocalisations that are response QTLs. (**d**) Colocalisation between a GWAS hit for rheumatoid arthritis (RA) and IFNγ-specific tuQTL at the *CD40* locus. PP4 represents the posterior probability from coloc [38] that the GWAS and QTL signals share a single causal variant. (**e**) Top panel: The lead GWAS variant (rs4239702) is associated with increased expression of the short 5′ UTR of the *CD40* gene. Bottom panel: Ensembl annotations couple the short 5′ UTR to skipped exon 6, but this is not supported by RNA-seq data (Figure S14). FPM, fragments per million. (**f)** Relative expression of the short 5′ UTR stratified by the genotype of the lead GWAS variant.

Finally, we assessed the condition-specificity of QTLs that colocalised with complex trait loci. We found that, on average, 12% of the GWAS colocalisations corresponded to response QTLs (Fig. 4c). One example is an IFNγ-specific promoter usage QTL for the *CD40* gene that colocalises with a GWAS signal for rheumatoid arthritis [36]. The alternative C allele of the rs4239702 variant is associated with increased usage of the transcript with the short 5′ UTR (Fig. 4e, f). This tuQTL was also visible at the absolute expression level of the two alternative promoters (Figure S14), but was missed by Leafcutter, because there is no change in junction reads. Although the variant was not significantly associated with total gene expression level (Figure S14), the two promoters contain the same start codon. As a result, the likely functional consequence of the *CD40* tuQTL is modulation of protein abundance. Although the same tuQTL was also detected at the full-length transcript usage level, the affected transcripts also differ from each other by alternatively spliced exon 6, making it challenging to interpret the result (Fig. 4e). The preferential upregulation of the transcript with the short 5′ UTR after exposure to an inflammatory stimulus is also supported by FANTOM5 capped analysis of gene expression (CAGE) data from primary macrophages (Figure S15) [37].

## Discussion

We have performed a comprehensive analysis of the genetic determinants of transcript usage in human iPSC-derived macrophages exposed to four different stimuli. Our approach to stratify transcripts into individual events greatly improved the interpretability of molecular mechanisms underlying tuQTLs. Consequently, we were able to discover that 55% of the transcript-level associations affected promoter or 3′ end usage and these variants were enriched in markedly different genomic features relative to canonical splicing QTLs. We also found that promoter usage QTLs were 50% more likely to be condition-specific than other transcriptional events and often colocalised with GWAS hits for complex traits. Event-level quantification also enabled us to assess the prevalence of transcriptional coupling between promoter usage, splicing and 3′ end usage. While a number of studies relying on full-length transcript quantification have suggested widespread coupling affecting majority of the genes [11,39], many of these could be false positives caused by missing transcripts (Fig. 1b). In contrast, both our event-level analysis as well as a recent full-length mRNA sequencing study [30] suggest that such coupling is relatively rare, affecting only ∼10% of the transcriptional events. This is further supported by a recent RNA-seq study of neural differentiation, which found that genes undergoing changes in promoter usage, alternative splicing or 3′ end usage are independent from each other [40]. Thus, event-level analysis might be preferable over transcript-level analysis when the aim is to identify specific transcriptional changes underlying genetic associations.

Choosing the optimal quantification method for RNA-seq data is a challenging problem. The field of detecting and quantifying individual transcriptional changes from RNA-seq data has been developing rapidly. One of the most successful approach has been the use of reads spanning exon-exon junctions to detect differential usage of individual exons within genes. In our study we used Leafcutter to perform junction-level analysis, but other options are available such as JUM [41] or MAJIQ [42]. A key advantage of junction-level analysis is that it can discover novel exon-exon junctions and is thus well-suited for characterising rare or unannotated splicing events. On the other hand, changes in 5′ and 3′ UTR length are not captured by junction-level methods, because these events do not overlap exon-exon junctions. Changes in UTR length can only be detected by methods that consider all reads originating from alternative transcript ends such as MISO [26] or txrevise proposed here. MISO provides more fine-grained events that can differentiate between various types of splicing events. Txrevise, on the other hand, provides a more comprehensive catalog of promoter and 3′ end events that can be continuously updated as reference annotations improve. A promising alternative to both of these methods is Whippet, which quantifies transcriptional events by aligning reads directly to the splice graph of the gene [43]. Thus, no single approach is consistently superior to others and characterizing the full spectrum of transcriptional consequences of genetic variation requires a combination of analytical strategies [16,25].

An important limitation of txrevise is that it is only able to quantify splicing events present in reference transcript databases. However, our approach can easily be extended by incorporating additional annotations such experimentally determined promoters from the FANTOM5 [44] projects or alternative polyadenylation sites from the PolyAsite database [31], as is done by QAPA [40]. Another option might be to incorporate novel transcripts identified by transcript assembly methods such as StringTie [45] into existing annotation databases. Nevertheless, since txrevise relies on Salmon for event-level quantification, it is still susceptible to some of the same limitations as full-length transcript quantification. Even though event-level analysis reduces the problem a bit, a positive transcript expression estimate does not guarantee that any specific exon is actually present in the transcript, especially if the transcript annotations are incomplete (Fig. 1b) [22]. Secondly, large eQTL effects and positional biases in the RNA-seq data can occasionally lead to spurious changes in transcript usage (Figures S9 and S10). Therefore, it is important to visually confirm candidate transcriptional events using either a genome browser or tools such as wiggleplotr [46] before embarking on follow-up experiments.

A key aim of QTL mapping studies is to elucidate the molecular mechanisms underlying complex trait associations. In our analysis, we found that over 50% of the genetic effects that colocalise with complex traits regulated transcript usage and did not manifest at the total gene expression level. Moreover, 42% of the transcript-level colocalisations affected promoter or 3′ end usage instead of splicing of internal exons. Importantly, no single quantification method was able to capture the full range of genetic effects, confirming that different quantification approaches often identify complementary sets of QTLs [16,25]. Consequently, there is great potential to discover additional disease associations by re-analysing large published RNA-seq datasets such as GTEx [47] with state-of-the-art quantification methods.

## Methods

### Cell culture and reagents

#### Donors and cell lines

Human induced pluripotent stem cells (iPSCs) from 123 healthy donors (72 female and 51 male) (Table S1) were obtained from the HipSci project [48]. Of these lines, 57 were initially grown in feeder-dependent medium and 66 were grown in feeder-free E8 medium. The cell lines were screened for mycoplasma by the HipSci project [48]. All samples for the HipSci resource were collected from consented research volunteers recruited from the NIHR Cambridge BioResource (http://www.cambridgebioresource.org.uk). Samples were collected initially under ethics for iPSC derivation (REC Ref: 09/H0304/77, V2 04/01/2013), with later samples collected under a revised consent (REC Ref: 09/H0304/77, V3 15/03/2013).

The details of the iPSC culture, macrophage differentiation and stimulation for the IFNγ + *Salmonella* study have been described previously [3] (Table S2). Macrophages for the acLDL study were obtained from the same differentiation experiments described above.

#### AcLDL stimulation

Macrophages differentiated from a total of 71 iPSC lines were used for the acLDL stimulation. Macrophages were grown in RPMI 1640 (Gibco) supplemented with 10% FBS (labtech), 2 mM L-glutamine (Sigma) and 100 ng/ml hM-CSF (R&D) at a cell density of 150,000 cells per well on a 6-well plate. On day 6 of the macrophage differentiation, two wells of the 6-well plate were exposed to 100 µg/ml human acLDL (Life Technologies) for 24 hours, whereas the other two wells were incubated in fresh RPMI 1640 medium without stimulation throughout this period.

For RNA extraction, cells were washed once with PBS and lysed in 300 µl of RLT buffer (Qiagen) per well of a 6-well plate. Lysates from two wells were immediately pooled and stored at −80°C. RNA was extracted using a RNA Mini Kit (Qiagen) following the manufacturer’s instructions and eluted in 35 µl nuclease-free water. RNA concentration was measured using NanoDrop, and RNA integrity was measured on Agilent 2100 Bioanalyzer using a RNA 6000 Nano Total RNA Kit.

### RNA sequencing and quality control

All RNA-seq libraries from the acLDL study were constructed manually using poly-A selection and the Illumina TruSeq stranded library preparation kit. The TruSeq libraries were quantified using Bioanalyzer and manually pooled for sequencing. The samples were sequenced on Illumina HiSeq 2000 using V4 chemistry and multiplexed at 6 samples/lane. The control and acLDL stimulated RNA samples from a single donor were always sequenced in the same experimental batch. Sample metadata is presented in Table S3. RNA-seq reads were aligned to the GRCh38 reference genome and Ensembl 87 transcript annotations using STAR v2.4.0j [49]. Subsequently, VerifyBamID v1.1.2 [50] was used to detect and correct any sample swaps between donors. Two samples from one donor (HPSI0513i-xegx_2) were excluded from downstream analysis, because they appeared to be outliers on the principal component analysis (PCA) plot of the samples.

### Quantifying gene and transcript expression

We used four alternative strategies to quantify transcription from RNA-seq data: (i) gene-level read count quantified with featureCounts [27], (ii) full-length transcript usage quantified with Salmon [28] (Fig. 1c), (iii) promoter, internal exon and 3′ end usage quantified with txrevise, and (iv) exon-exon junction usage quantified with Leafcutter [6].

#### Gene-level read count

We used featureCounts v1.5.0 [27] to count the number of uniquely mapping fragments overlapping transcript annotations from Ensembl 87. We excluded short RNAs and pseudogenes from the analysis leaving 35,033 unique genes of which 19,796 were protein coding. Furthermore, in both IFNγ + *Salmonella and* acLDL dataset we used only genes with mean expression in at least one of the conditions greater than 1 transcripts per million (TPM) [51] in all downstream analyses. This resulted in 12,660 and 12,103 genes included for analysis in the IFNγ + *Salmonella and* acLDL datasets, respectively. We quantile-normalised the data and corrected for sample-specific GC content bias using the conditional quantile normalisation (cqn) [52] R package as recommended previously [53].

#### Full-length transcript usage

We downloaded the FASTA files with messenger RNA (mRNA) and non-coding RNA sequences from the Ensembl website (version 87). We concatenated the two files and used salmon v0.8.2 [28] with ‘--seqBias --gcBias --libType’ options to quantify the expression level of each transcript. We used tximport [54] package to import the expression estimates into R and calculated the relative expression of each transcript by dividing the TPM expression estimate of each transcript with the sum of the expression estimates of all transcripts of the gene.

#### Quantifying transcriptional events with txrevise

We downloaded exon coordinates for all Ensembl 87 transcripts using the `makeTxDbFromBiomart` function from the GenomicFeatures [55] R package. We also downloaded metadata for these transcripts using the biomart [56] R package. Finally, we extracted transcript tags from the GTF file downloaded from the Ensembl website. This step was necessary, because Ensembl contains a large number of truncated transcripts (marked with cds_start_NF or cds_end_NF tags) (Figure S1), but this information is not present in biomart.

We developed the txrevise R package to pre-process transcript annotations prior to quantification. First, we extended all truncated protein coding transcripts using exons from the longest annotated transcript of the gene that was part of the GENCODE Basic gene set (Figure S2). We also performed the same step on transcripts annotated in Ensembl as `retained_intron, processed_transcript or nonsense_mediated_decay,` because they often ended abruptly in the middle of the exons and were unlikely to correspond to true transcription start and end sites.

Next, we focused on splitting full-length transcripts into alternative promoters, internal exons and 3′ ends. However, some genes contained either non-overlapping transcripts or very short transcripts that complicated this process. Thus, for each gene we first identified a subset of transcripts that shared the largest number of exons with each other. We then used the shared exons as a scaffold and constructed sets of independent promoters, internal exons and 3′ ends (group 1) (Figure S3). We repeated this process for a second subset of transcripts that shared the most exons with each other (group 2) (Figure S3). Thus, the original transcripts from each gene were split into up to six sets of transcriptional events (two groups of alternative promoters, internal exons and 3′ ends). Next, to ensure that the new alternative promoter and 3′ end events did not capture splicing changes, we masked all alternative exons that were not the first or last exons (Figure S4). This final step can optionally be skipped to discover more association at the expense of losing some interpretability, because a subset of the promoter and 3′ end events might be tagging splicing changes. We used Salmon [28] to independently quantify the expression of each set of transcriptional events. Finally, we used tximport [54] to import the expression estimates into R and calculated the relative expression of each transcriptional event by dividing the TPM expression estimate of each event with the sum of the expression estimates of all events in one group.

#### Quantifying intron excision ratios with Leafcutter

Finally, we used Leafcutter [6] to quantify the relative excision frequencies of alternative introns. We used the spliced alignments from STAR as input to Leafcutter. We did not correct for reference mapping bias, because we wanted to be able to directly compare Leafcutter results with those from Salmon and there is no obvious way to correct for reference mapping bias in Salmon quantification. We used the default parameters of requiring at least 50 reads supporting each intron cluster and allowing introns of up to 500 kb in length.

### Mapping expression and transcript usage QTLs

#### Preparing genotype data

We obtained imputed genotypes for all of the samples from the HipSci [48] project. We used CrossMap v0.1.8 [57] to convert variant coordinates from GRCh37 reference genome to GRCh38. Subsequently, we filtered the VCF file with bcftools v.1.2 to retain only bi-allelic variants (both SNPs and indels) with IMP2 score > 0.4 and minor allele frequency (MAF) > 0.05. We created a separate VCF files for the IFNγ + *Salmonella* study (84 individuals) and the acLDL study (70 individuals). The same VCF files were used for all downstream analyses and were imported into R using the SNPRelate [58] R package.

#### Association testing

We used QTLTools [59] to map QTLs in two stages. First, we used the permutation pass with ‘‘--permute 10000 --grp-best’ options to calculate the minimal lead variant p-value for each feature (gene, transcript or splicing event) in a +/-100 kb window around each feature. The ‘--grp-best’ option ensured that in case of transcript usage QTLs, the permutation p-values were corrected for the number of alternative transcripts, exon-exon junction or transcriptional events tested. Secondly, we used the nominal pass to calculate nominal association p-values in a +/-500 kb *cis* window around each feature. We used a larger *cis* window for the nominal pass to ensure that we did not have missing data in the colocalisation analysis (see below), where we used the +/-200 kb *cis* window around each lead QTL variant. We included the first six principal components of the phenotype matrix as covariates in the QTL analysis.

### QTL replication between quantification methods

To compare the QTLs detected by different quantification methods, we estimated the fraction of QTL lead variants detected by each method that were replicated by the other methods. Since read count and full-length transcript usage analysis were performed at the gene level, we decided to perform the replication analysis at the gene level as well. Because txrevise and leafcutter quantified multiple events per gene and sometimes detected multiple independent QTLs (Figure S6), we picked the lead variant with the smallest p-value across all of the events quantified for a given gene as the gene-level lead variant. For each pairwise comparison of quantification methods, we first identified all lead variant-gene pairs with FDR < 0.01 detected by the query method. Subsequently, we extracted the lead variants for the same genes detected by the replication method and estimated the fraction of those that were in high LD (R^2^ < 0.8) with each other. We then repeated this analysis for all pairs of quantification methods. Note that this measure is not necessarily symmetric between the quantification methods and also depends on the statistical power of each method. Since Leafcutter was less powered than other methods, it also replicated smaller fraction of QTLs detected by the other methods. In contrast, 50% of the Leafcutter QTLs were replicated by txrevise and full-length transcript usage (Fig.2b).

### QTL enrichment in genomics annotations

#### Constructing genomic annotations

##### Gene features

We downloaded transcript annotations from Ensembl version 87 [23] using the GenomicFeatures [55] R package. We retained only protein coding transcripts and used `fiveUTRsByTranscript, threeUTRsByTranscript, cdsBy, intronsByTranscript` and promoters functions to extract 5′ UTRs, 3′ UTRs, coding sequences, introns and promoters, respectively. We defined promoters as sequences 2000 bp upstream and 200 bp downstream of the annotated transcription start sites.

##### Polyadenylation sites

We downloaded the coordinates of experimentally determined human polyadenylation sites from the PolyASite database (version r1.0) [31]. After converting the coordinates to the GRCh38 reference genome with CrossMap [57], we extended each polyadenylation site to +/-25 bp from the center of the site.

##### Chromatin accessibility

We downloaded the coordinates of accessible chromatin regions in macrophages across four conditions (N, I, S, I + S) from our previous study [3]. Specifically, we downloaded the ATAC_peak_metadata.txt.gz file from Zenodo (https://doi.org/10.5281/zenodo.1170560).

##### RNA binding proteins

We downloaded processed eCLIP [60] peak calls for 93 RNA binding proteins (RBPs) [32] from the ENCODE web site (https://www.encodeproject.org). Each protein was measured in two biological replicates, resulting in 186 sets of peaks. We only used data from the K562 myelogenous leukemia cell line. We further used Supplementary Table 1 from [32] to identify a subset of 29 RBPs that have previously been implicated in splicing regulation, five factors that have been implicated in 3′ end processing and two factors (SRSF7 and HNRNPK) that have been implicated in both. Within each group (splicing, 3′ end processing and both), we first removed all peaks that were detected only once and then merged all peaks into a single genomic annotation.

#### Enrichment analysis

We used fgwas v0.3.6 [33] with the ‘-fine’ option to identify the genomic annotations in which different types of QTLs were enriched. We converted QTLtools p-values to z-scores using the `stats.norm.ppf(p/2, loc=0, scale=1)` function from SciPy [61], where p is the p-value from QTLtools. The sign of the z-score was determined based on the sign of the QTL effect size. We included all genomic annotations into a joint fgwas model using the ‘-w’ option. For the enrichment analysis we used QTLs from the naive condition only, but we found that the enrichments patterns were very similar in all stimulated conditions.

### Overlap with genome-wide association studies

#### Summary statistics

We obtained full summary statistics for ten immune-mediated disorders: inflammatory bowel disease (IBD) including ulcerative colitis (UC) and Crohn’s disease (CD) [62], Alzheimer’s disease (AD) [63], rheumatoid arthritis (RA) [36], systemic lupus erythematosus (SLE) [64], type 1 diabetes (T1D) [65], schizophrenia (SCZ) [66], multiple sclerosis (MS) [67], celiac disease (CEL) [68] and narcolepsy (NAR) [69]. We also obtained summary statistics for type 2 diabetes (T2D) [70], cardiovascular disease (CAD) [71,72] and myocardial infarction (MI) [71]. Finally, we obtained summary statistics for 20 cardiometabolic traits from a recent meta-analysis [73]. Summary statistics for T1D, CEL, IBD, RA, AD, MS and SLE were downloaded in 2015. SCZ, T2D and NAR were downloaded in 2016. T2D summary statistics were converted from GRCh36 to GRCh37 coordinates using the LiftOver tool, all of the other summary statistics already used GRCh37 coordinates.

#### Colocalisation analysis

We used coloc v2.3-1 [38] to test for colocalisation between gene expression and transcript usage QTLs and GWAS hits. We ran coloc on a 400 kb region centered on each lead eQTL and tuQTL variant that was less than 100 kb away from at least one GWAS variant with a nominal p-value < 10^-5^. We used the following prior probabilities: p1 = 10^-4^, p2 = 10^-4^ and p12 = 10^-5^. We then applied a set of filtering steps to identify a stringent set of eQTLs and tuQTLs that colocalised with GWAS hits. Similarly to a previous study [74], we first removed all cases where PP3 + PP4 < 0.8, to exclude loci where we were underpowered to detect colocalisation. We then required PP4/(PP3+PP4) > 0.9 to only keep loci where coloc strongly preferred the model of a single shared causal variant driving both association signals over a model of two distinct causal variants. We excluded all colocalisation results from the MHC region (GRCh38: 6:28,510,120-33,480,577) because they were likely to be false positives due to complicated LD patterns in this region. We only kept results where the minimal GWAS p-value was < 10^-6^. Plots illustrating the sharing of colocalised GWAS signals by different quantification methods were made using UpSetR [75].

### Code availability

The Snakemake [76] files used for gene and transcript expression quantification, QTL mapping and colocalisaton are available from the project’s GitHub repository (https://github.com/kauralasoo/macrophage-tuQTLs). The same repository also contains R scripts that were used for all data analysis and figures. The txrevise R package is available from GitHub (https://github.com/kauralasoo/txrevise) and wiggleplotr R package that was used to make transcript read coverage plots is available from Bioconductor (http://bioconductor.org/packages/wiggleplotr/).

## Data availability

RNA-seq data from the acLDL stimulation study is available from ENA (ERP022909) and EGA (EGAS00001000876). RNA-seq data from the IFNγ + *Salmonella* study is available from ENA (ERP020977) and EGA (EGAS00001002236). The imputed genotype data for HipSci cell lines is available from ENA (ERP013161) and EGA (EGAD00010000773). Processed data and QTL summary statistics are available from Zenodo: https://zenodo.org/communities/macrophage-tuqtls/.

## Acknowledgements

We thank Jeremy Schwartzentruber and Leopold Parts for their helpful comments on the manuscript. We thank WTSI DNA Pipelines and Cytometry Core Facility for their sequencing and flow cytometry services. We thank Elena Vigorito and Joanna M.M Howson (Cardiovascular Epidemiology Unit) for their help with statistical analyses during the early phases of this project. This work was supported by the Wellcome Trust (WT098051) and the British Heart Foundation Cambridge Centre of Excellence (RE/13/6/30180). K.A. was supported by a PhD fellowship from the Wellcome Trust (WT099754/Z/12/Z), a postdoctoral fellowship from the Estonian Research Council (MOBJD67) and a grant from the Estonian Research Council (IUT34-4). The Cardiovascular Epidemiology Unit is supported by the UK Medical Research Council (MR/L003120/1), British Heart Foundation (RG/13/13/30194) and NIHR Cambridge Biomedical Research Centre. The iPSC lines were generated at the Wellcome Sanger Institute, under the Human Induced Pluripotent Stem Cell Initiative funded by a strategic award (WT098503) from the Wellcome Trust and Medical Research Council. We also acknowledge Life Science Technologies Corporation as the provider of cytotune. This work was carried out in part at the High Performance Computing Center of University of Tartu.

## Author contributions

K.A., D.S.P. and D.J.G. wrote the paper with input from all authors. K.A. and J.R. performed the macrophage differentiation experiments. J.R. optimised and performed the acLDL stimulation experiments. K.A. analysed the data. K.A., D.S.P., D.F.F. and D.J.G. designed the experiments. D.S.P., D.J.G. and J.D. supervised the research.

## Supplementary Figures

**Figure S1.**
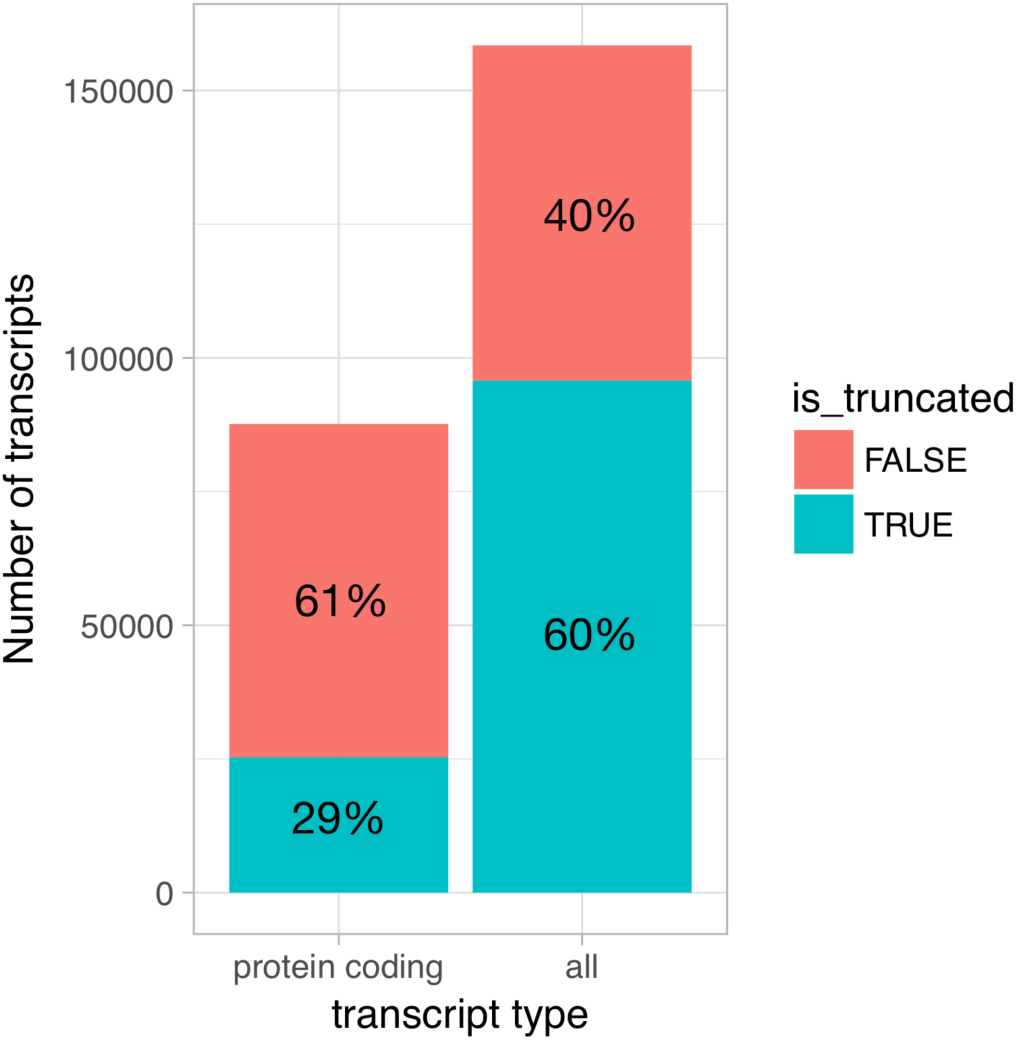
Truncated transcripts in the Ensembl database. For protein coding transcripts, we extracted the `cds_start_NF` and `cds_end_NF` fields from the Ensembl v87 GTF file to identify transcripts that were truncated at either 5′ or 3′ ends. For non-protein coding transcripts, we considered all transcripts annotated as `nonsense_mediated_decay, processed_transcript or retained_intron` to be truncated at both 5′ or 3′ ends, because we observed that many of those started and ended abruptly in the middle of exons. We included only protein coding genes in the analysis.

**Figure S2.**
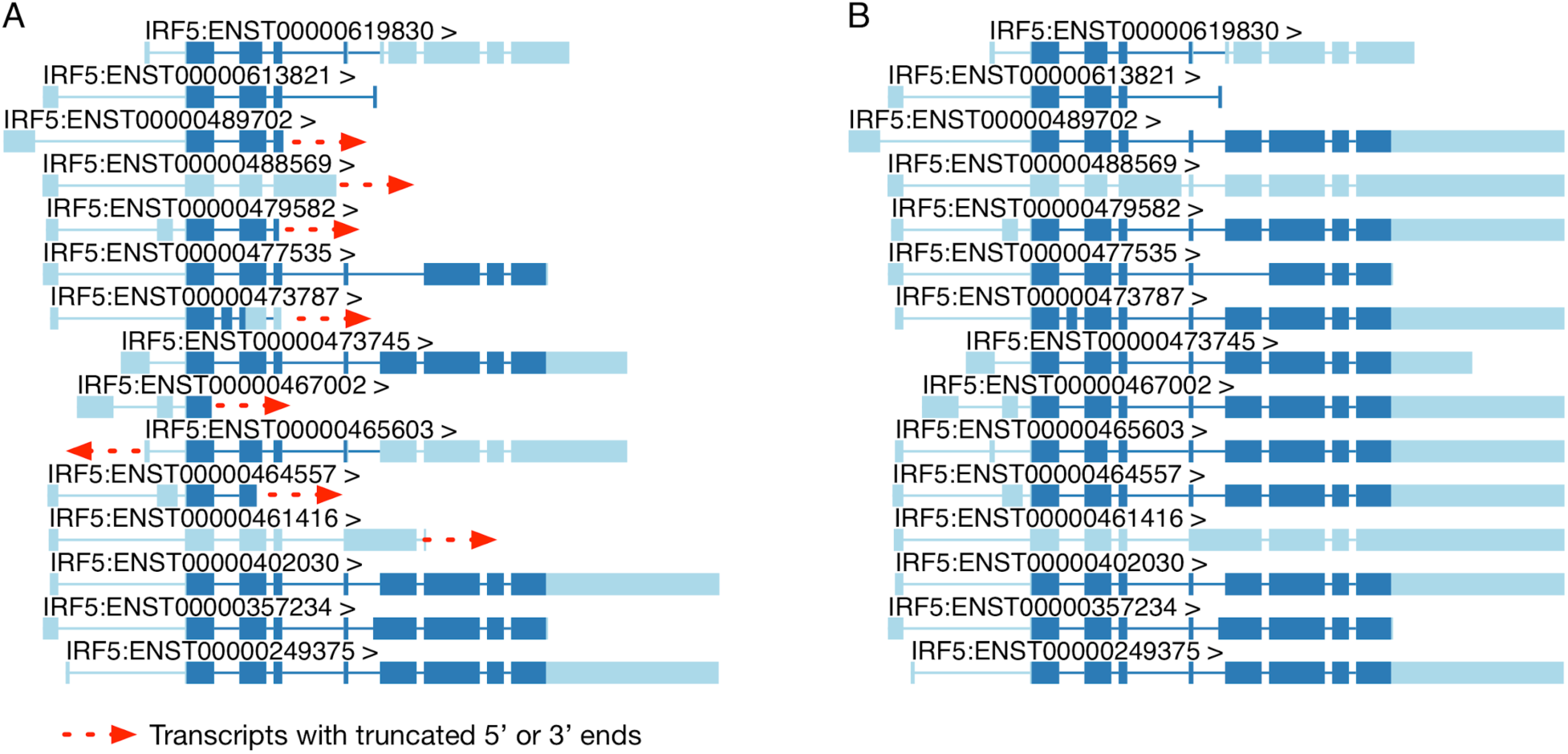
Extending truncated transcript annotations. (**A**) Original transcript annotations for *IRF5* in the Ensembl database. (**B**) Txrevise extends truncated *IRF5* transcripts by copying over exons from the longest transcript of the gene (ENST00000402030).

**Figure S3.**
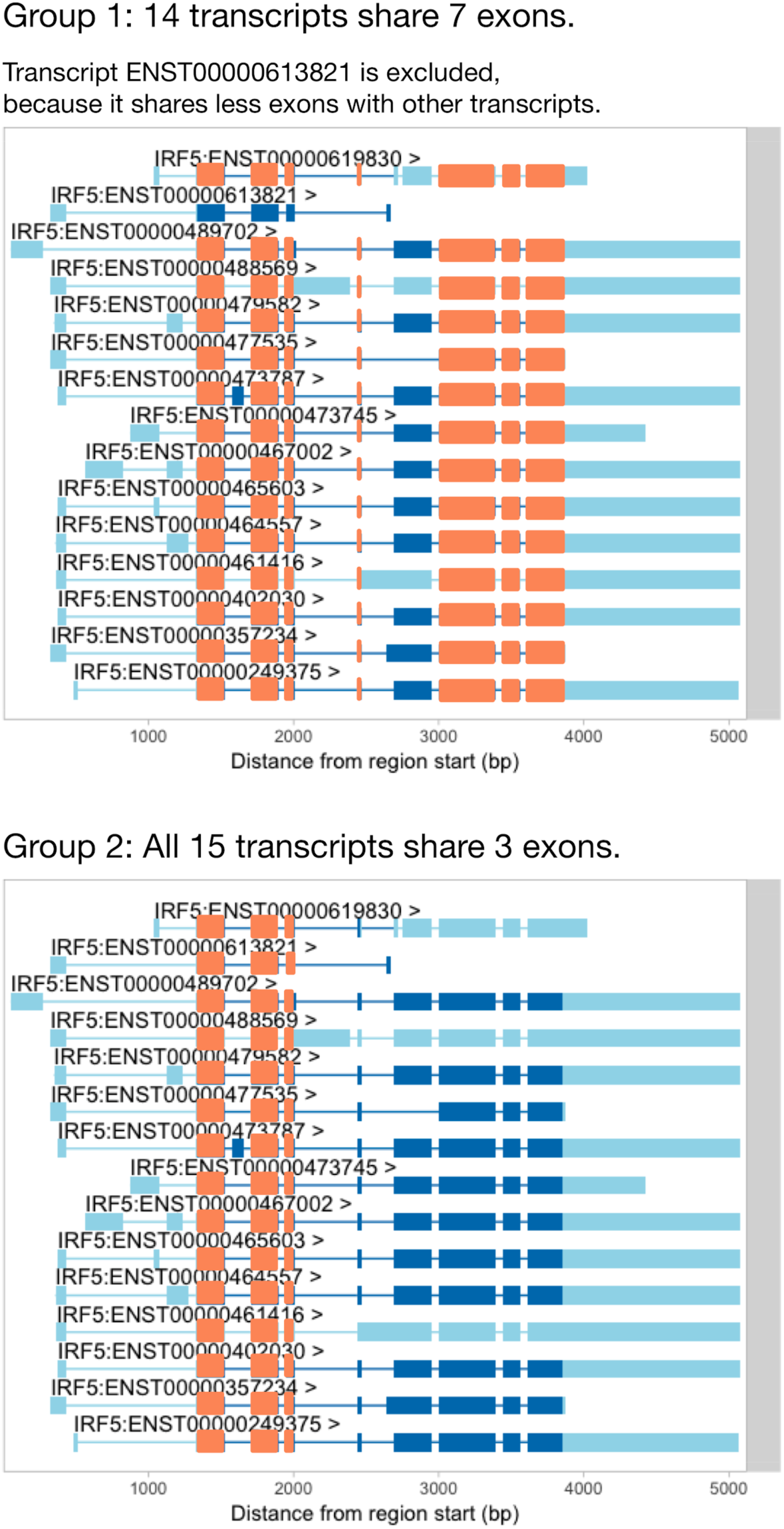
Before we can use txrevise to stratify transcripts into events, we need to identify a subset of transcripts that all share at least one exon. *IRF5* has three exons that are shared between all of the transcripts and so we could use those as a scaffold for txrevise to construct independent transcriptional events (group 2). However, some genes do not have any exons that are shared across all transcripts. In that case, it might be preferential to choose the largest subset of transcripts that share the most exons (group 1). Furthermore, even in the case of *IRF5*, one transcript (ENST00000613821) is much shorter than others and excluding it might lead to better stratification of transcripts into alternative promoter, internal exon and 3′ end events (group 1).

**Figure S4.**
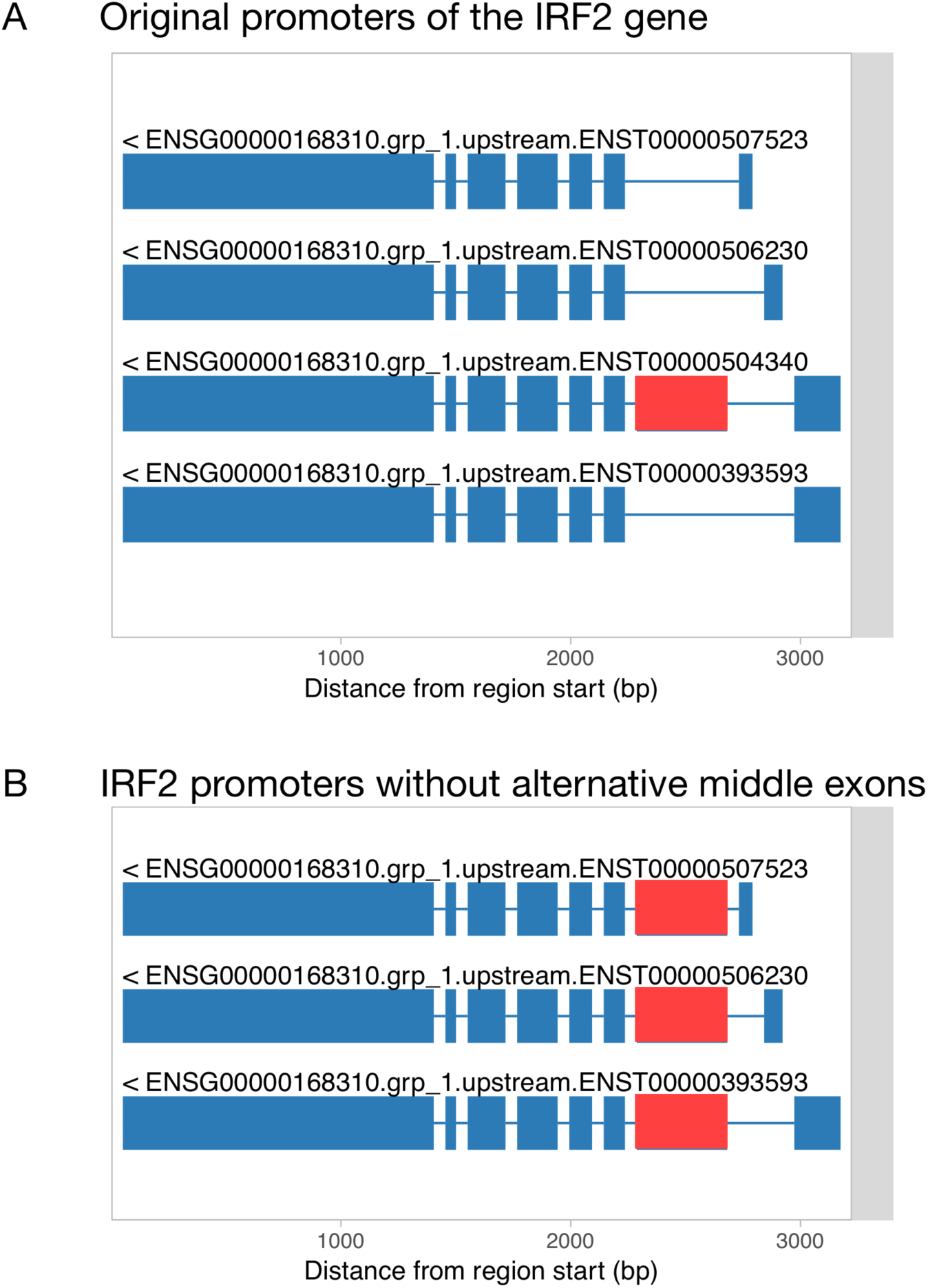
Filling in alternative internal exons for promoter and 3′ end events. (**A**) Original transcript start events constructed by txrevise contains an alternative second exon. (**B**) To construct promoter events, the alternative second exon is added into all transcripts and redundant transcripts are removed.

**Figure S5.**
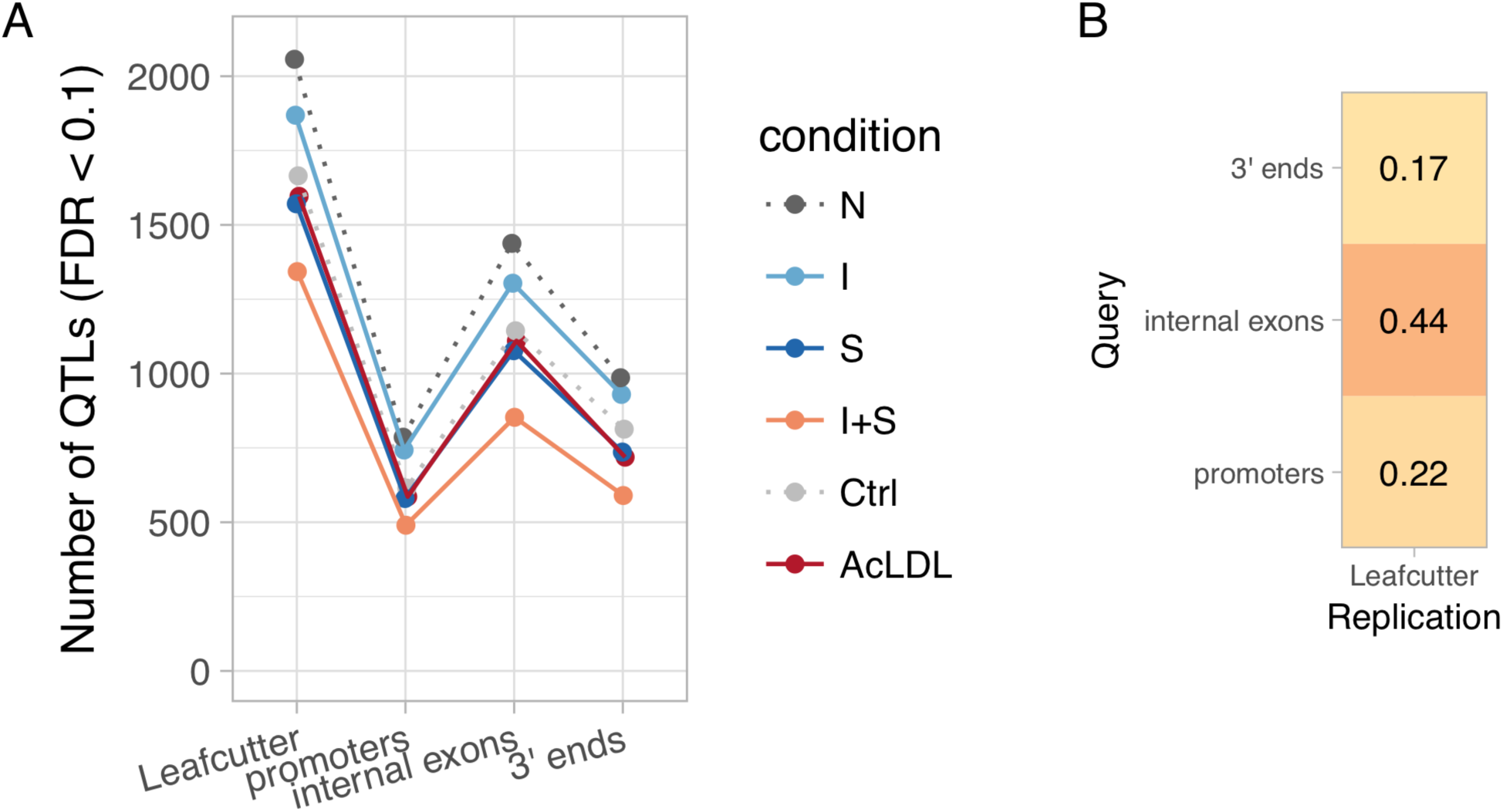
Diversity of transcript usage QTLs. (**A**) Number of detected transcript usage QTLs affecting exon-exon junction usage (Leafcutter) or different parts of the transcript (promoters, internal exons, 3′ ends) in each experimental condition. (**B**) Fraction of txrevise promoter, internal exon and 3′ end usage QTLs that were replicated by Leafcutter (lead variant within R^2^ > 0.8 for the same gene; see Methods).

**Figure S6.**
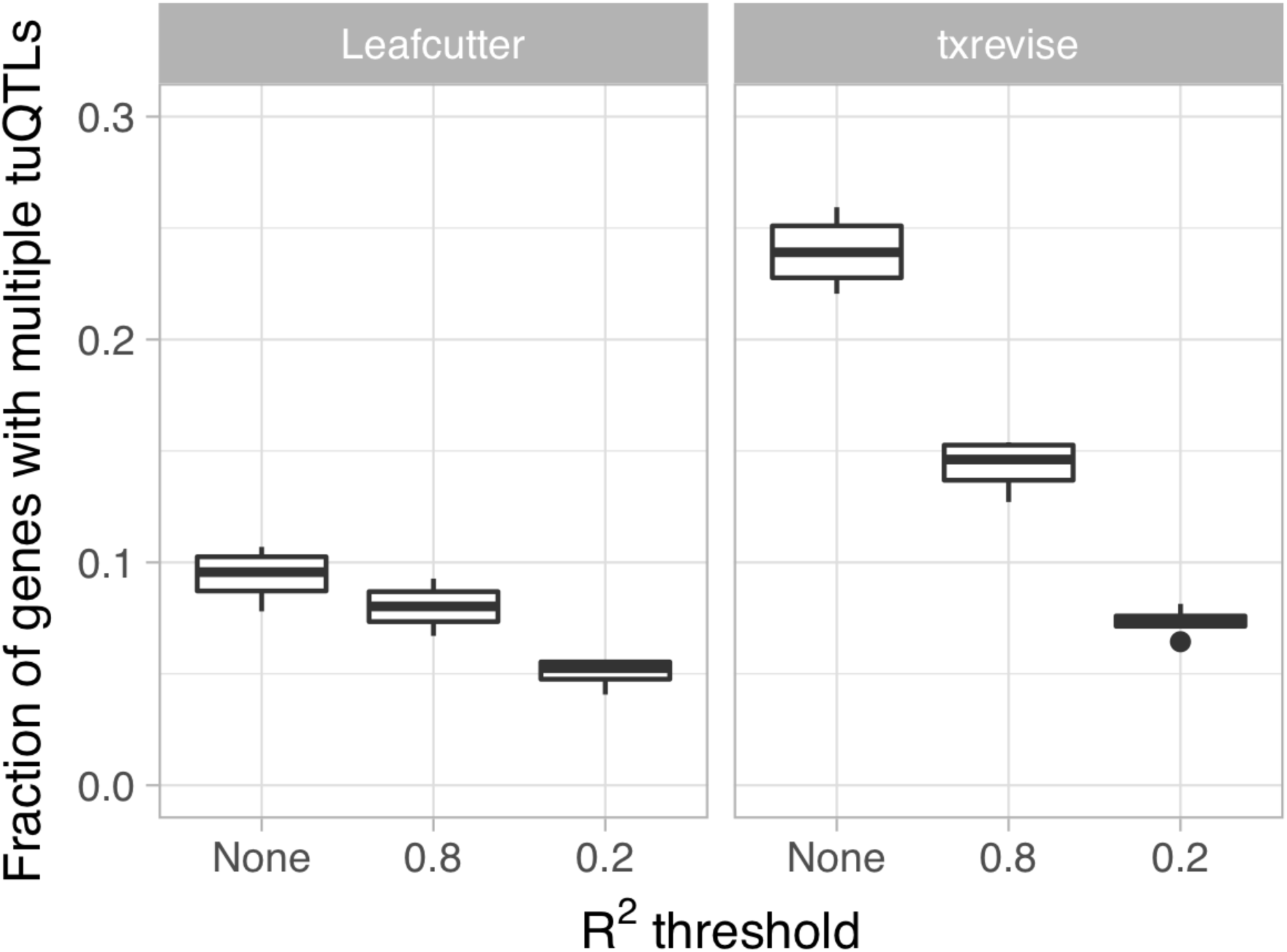
Fraction of genes with multiple independent tuQTLs detected by Leafcutter and txrevise. We first identified all tuQTLs for the same gene at the same 10% FDR threshold and then ascertained their independence by thresholding on LD at two levels of stringency (R^2^ < 0.2 or R^2^ < 0.8).

**Figure S7.**
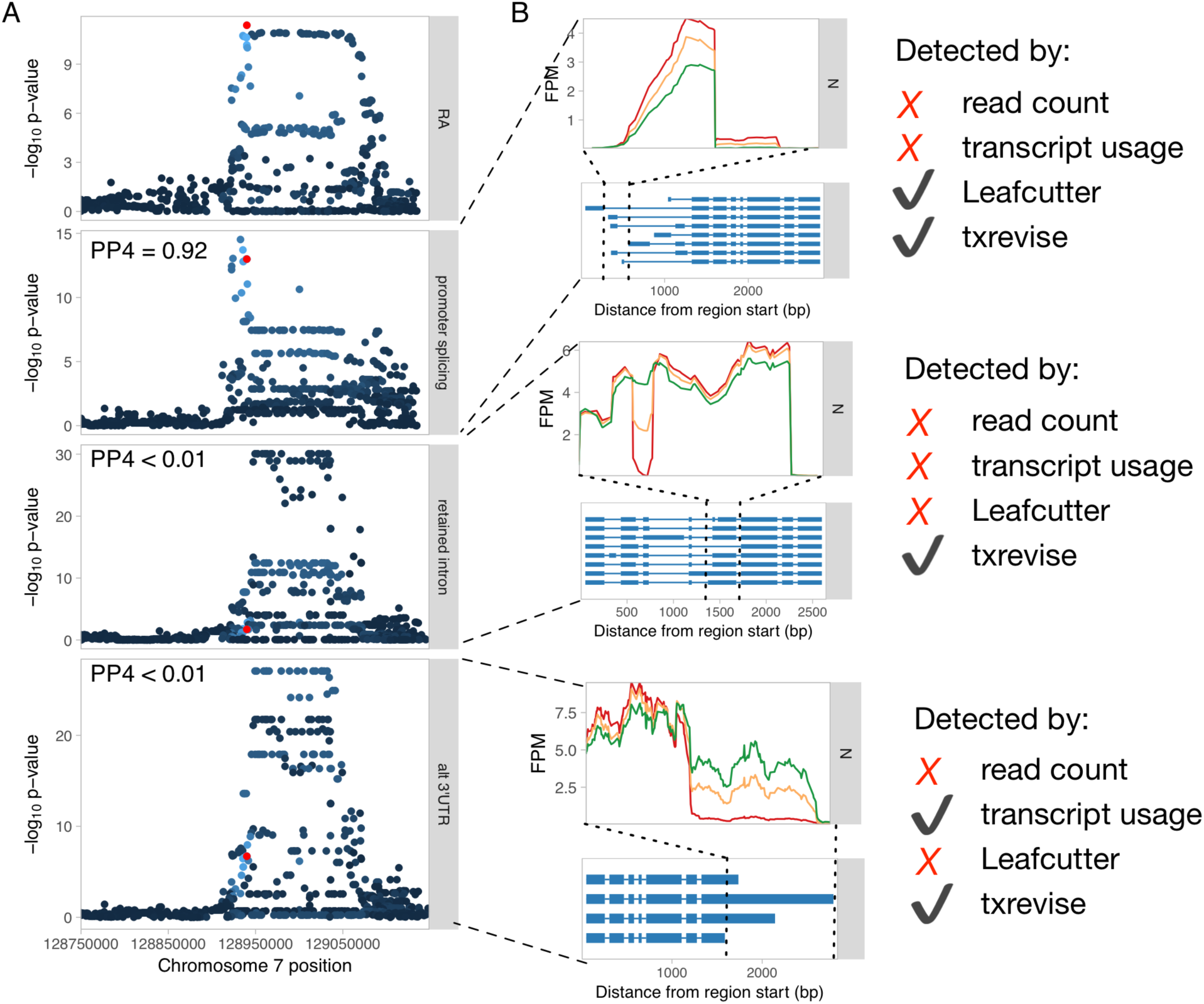
Genetics of transcript usage of the *IRF5* gene. (**A**) Three independent tuQTLs for *IRF5* regulating splicing in the first exon, intron retention in coding exon 5, and 3′ UTR length. PP4 represents the posterior probability from coloc [38] that the GWAS and QTL signals share a single causal variant. Only the promoter splicing QTL colocalises with a GWAS hit for rheumatoid arthritis (RA) [36] (PP4 = 0.92). (**B**) RNA-seq read coverage plots of the three tuQTLs stratified by the genotypes of the lead variants. Only txrevise detected all three tuQTLs. Leafcutter only detected the first splicing event, because the other two did not manifest at level of junction reads. Similarly, full-length transcript usage analysis only detected the polyadenylation event, because it had the largest effect size. FPM, fragments per million.

**Figure S8:**
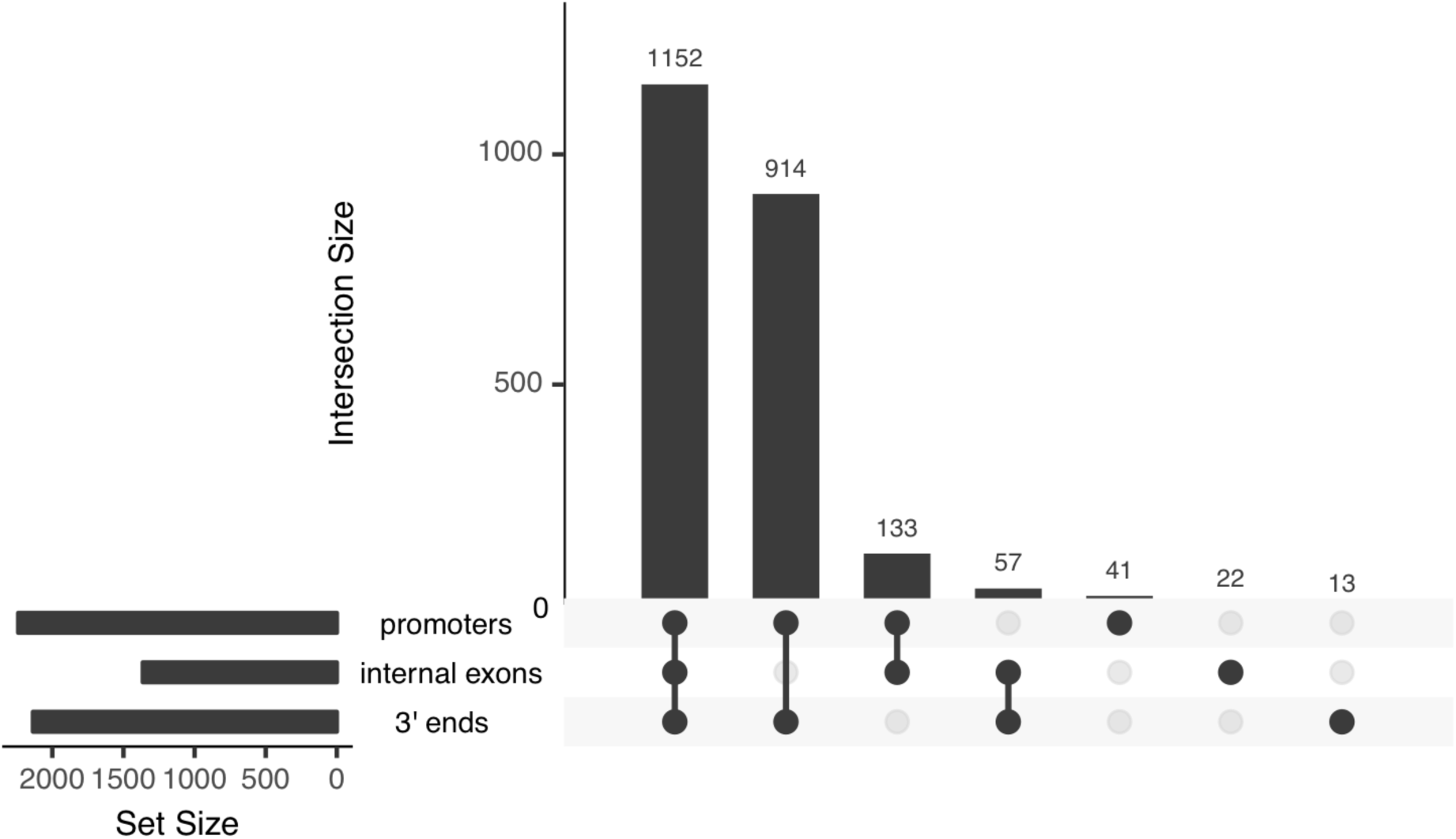
Naive estimates of transcriptional coupling between promoters, internal exons and 3′ ends based on full-length transcript quantification. Only 97/2332 (3.2%) tuQTLs appear to be associated with a single transcriptional event.

**Figure S9:**
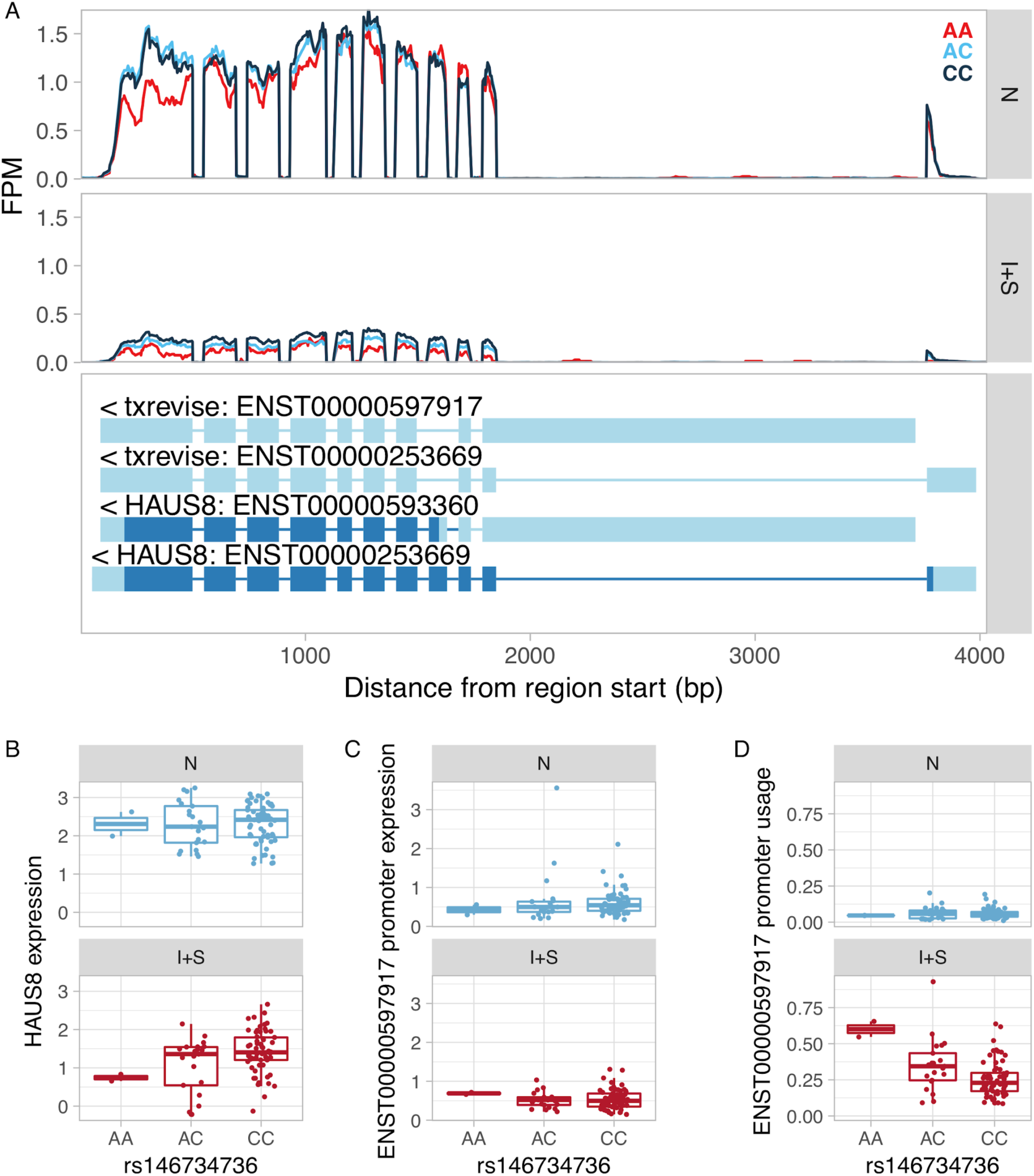
Example of an apparent tuQTL caused by an IFNγ + *Salmonella* specific eQTL. (**A**) RNA-seq read coverage plot of the *HAUS8* gene stratified by the genotype of the lead eQTL variant. FPM, fragments per million. (**B**) The rs146734736 variant is an IFNγ + Salmonella-specific eQTL for the *HAUS8* gene, regulating its total gene expression level. (**C**) The lead eQTL variant is not associated with the absolute expression of the alternative promoter of the gene (ENST00000597917). Furthermore, the average expression of the transcript with the alternative promoter is below 1 TPM, suggesting that it is either not expressed or expressed at a very low level. (**D**) Since the rs146734736 variant is associated with total expression level of the gene, but not with the absolute expression level of the ENST00000597917 transcript, it appears to be associated with the *relative* expression of the ENST00000597917 transcript.

**Figure S10.**
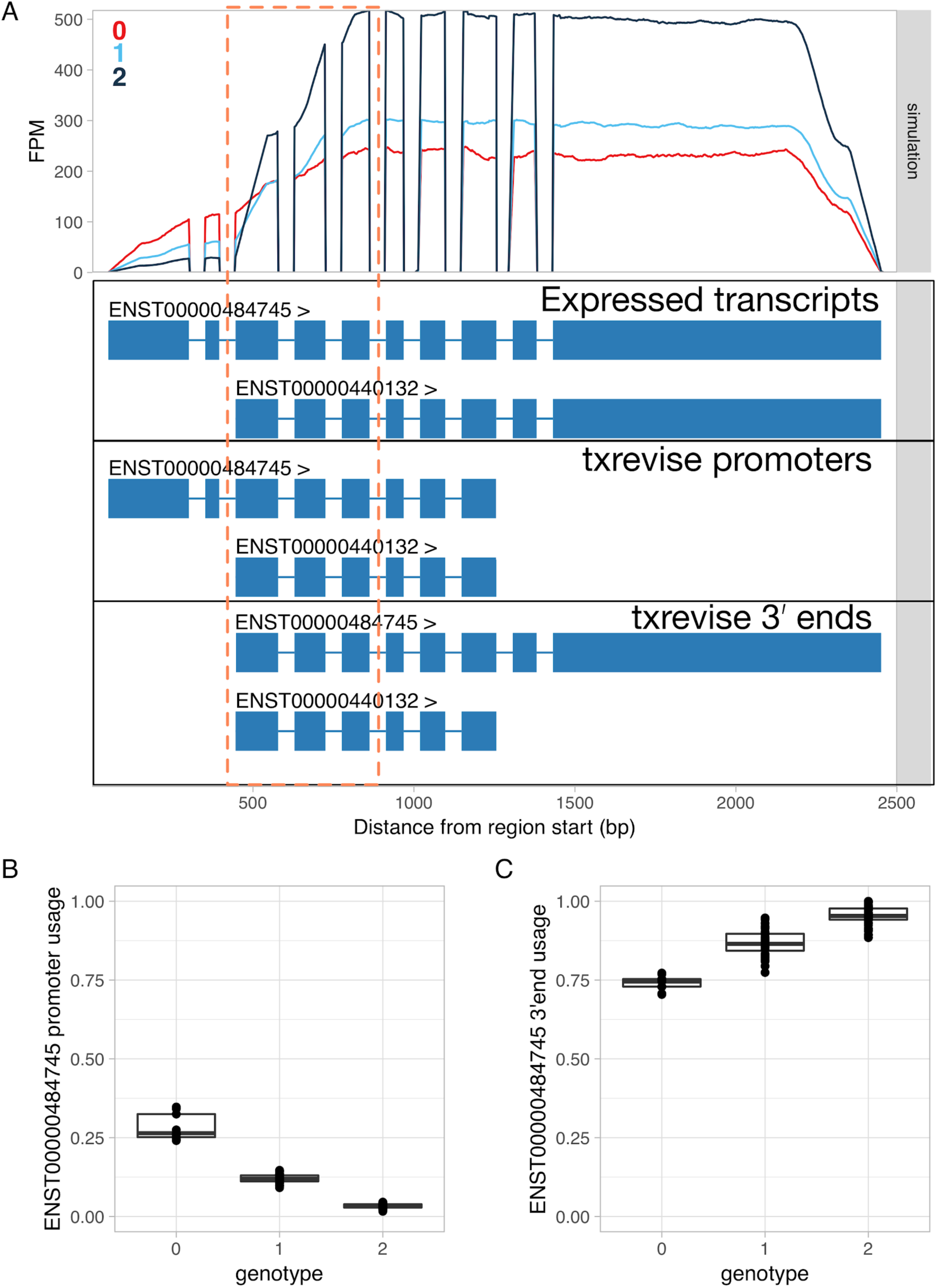
Simulated promoter usage QTL for the *RNF220* gene leads to a false positive association at the 3′ end. Simulations were performed using polyester [77]. (**A**) Simulated read coverage of the *RNF220* gene. Each additional copy of the alternative allele was simulated to decrease the usage of the long 5′ end over the short 5′ end (*Expressed transcripts*). Note that due to positional bias [78,79] in the RNA-seq data, the read coverage gradually decreases at the 5′ and 3′ ends of the transcript. The extent of this bias is likely to depend on the fragment length distribution of the dataset. While the simulated data had fragment length distribution with mean = 250 and sd = 25, this can vary substantially for real-world datasets. Importantly, increased usage of the short 5′ end leads to lower read coverage at exons 3-5 due to this bias (highlighted by the dashed orange box). FPM, fragments per million. (**B**) Estimated usage of the long 5′ end event stratified by the genotype of the simulated causal variant. In this case, txrevise is able to correctly detect the decrease in long 5′ end usage. (**C**) Estimated usage of the transcript with the long 3′ end stratified by the causal promoter usage QTL variant. Although we simulated no change at the 3′ end of the gene, txrevise still detects a false positive association. This is due to the fact that individuals with higher expression of the transcript with a short 5′ end have relatively fewer reads mapping to exons 3-5 (dashed orange box) compared to the exons that are specific to the long 3′ end. Consequently, it seems that the genetic variant decreasing short 5′ end usage is also associated with increased expression of the long 3′ end even though there are no additional reads mapping to the long 3′ end.

**Figure S11.**
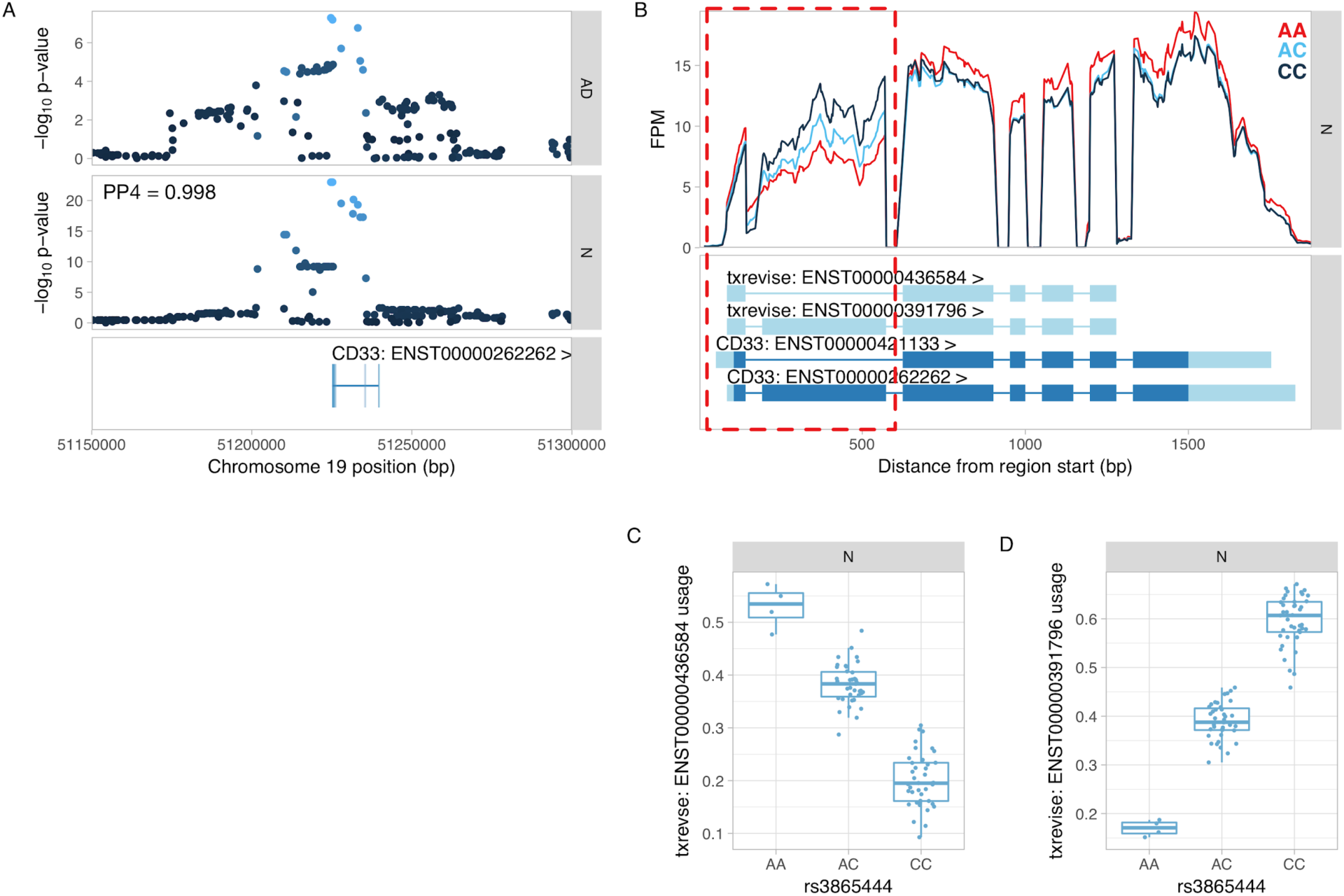
Colocalisation between *CD33* splicing QTL (sQTL) and GWAS hit for Alzheimer’s disease (AD). (**A**) Manhattan plots of the Alzheimer’s disease GWAS hit [63] and a sQTL for *CD33*. PP4 represents the posterior probability from coloc [38] that the GWAS and sQTL signals share a single causal variant. (**B**) Read coverage plot of the of the *CD33* gene stratified by the genotype of the lead sQTL variant (rs3865444). The alternatively spliced exon 2 is highlighted by the red rectangle. Ensembl transcript annotations falsely link skipped exon 2 to alternative 5′ and 3′ UTRs although these do not appear to be regulated by the sQTL variant. FPM, fragments per million. (**C**) Usage of the *CD33* transcript with skipped exon 2 stratified by the lead sQTL variant. (**D**) Usage of the *CD33* transcript containing exon 2 stratified by the lead sQTL variant.

**Figure S12.**
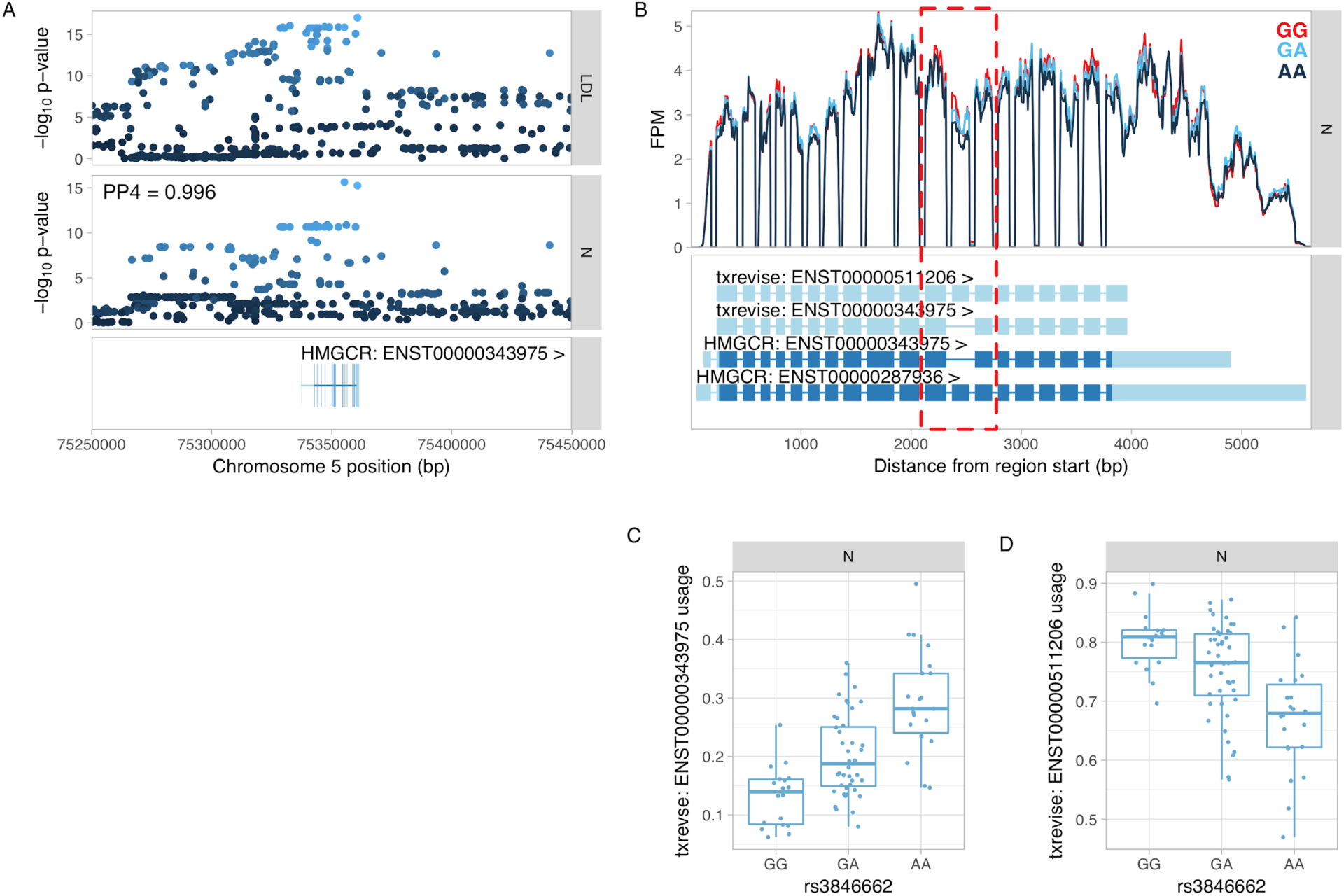
Colocalisation between *HMGCR* splicing QTL (sQTL) and GWAS hit for LDL. (**A**) Manhattan plots of the LDL GWAS hit [73] and an sQTL for *HMGCR*. PP4 represents the posterior probability from coloc [38] that the GWAS and sQTL signals share a single causal variant. (**B**) Read coverage plot of the of the *HMGCR* gene stratified by the genotype of the lead sQTL variant (rs3846662). The alternatively spliced exon 13 is highlighted by the red rectangle. Ensembl transcript annotations falsely link skipped exon 13 to alternative 5′ and 3′ UTRs although these do not appear to be differentially regulated by the sQTL variant. FPM, fragments per million. (**C**) Usage of the *HMGCR* transcript with skipped exon 13 stratified by the lead sQTL variant. (**D**) Usage of the *HMGCR* transcript containing exon 13 stratified by the lead sQTL variant.

**Figure S13.**
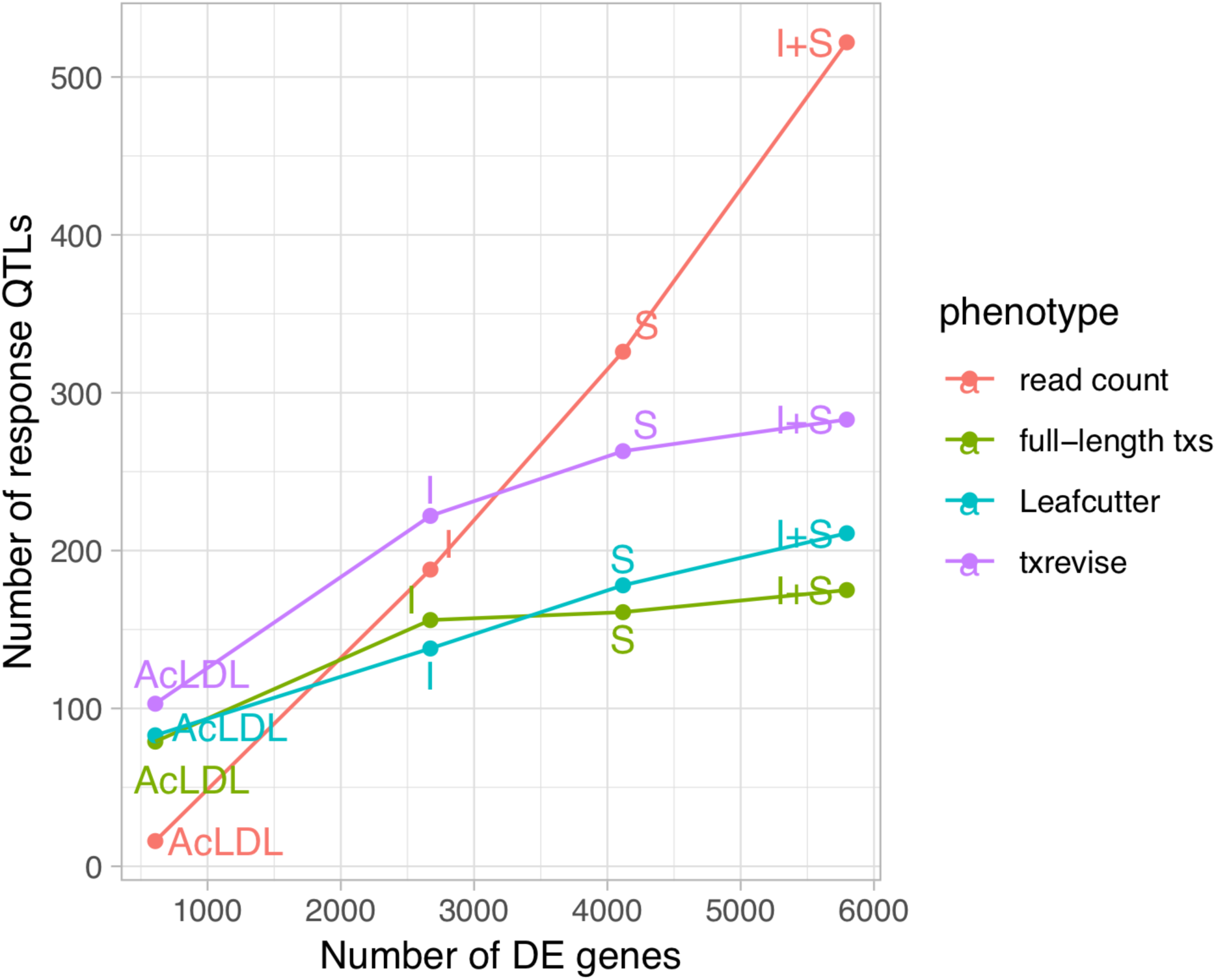
Relationship between the number of differential expressed genes in each condition and number of response QTLs detected in that condition.

**Figure S14.**
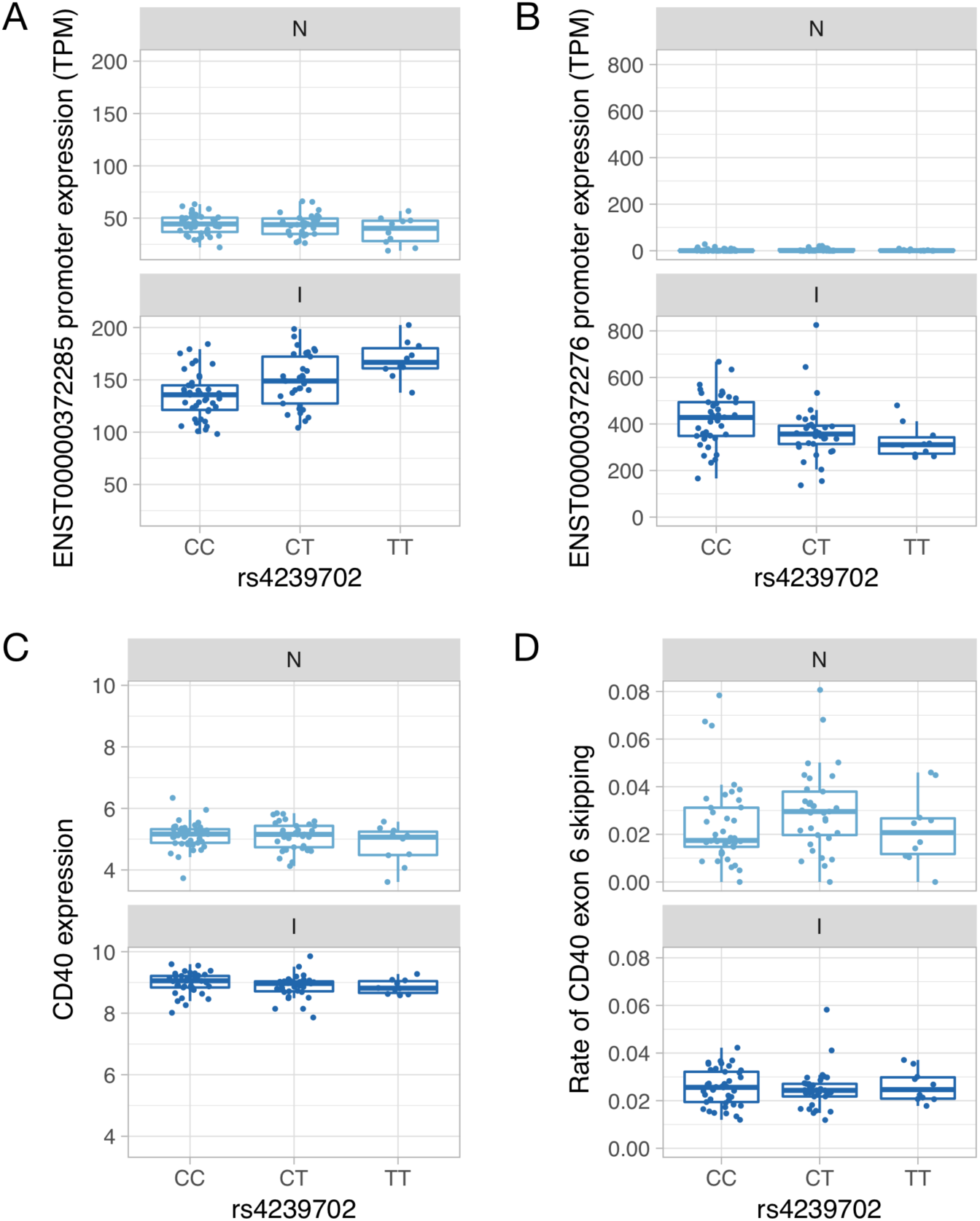
Genetics of *CD40* expression. (**A**) Absolute expression of the ENST00000372285 promoter in TPM units stratified by the genotype of the rs4239702 variant. (**B**) Absolute expression of the ENST00000372276 promoter in TPM units stratified by the genotype of the rs4239702 variant. (**C**) Normalized *CD40* read count stratified by the genotype of the rs4239702 variant. (**D**) Rate of *CD40* exon 6 skipping stratified by the genotype of the rs4239702 variant.

**Figure S15.**
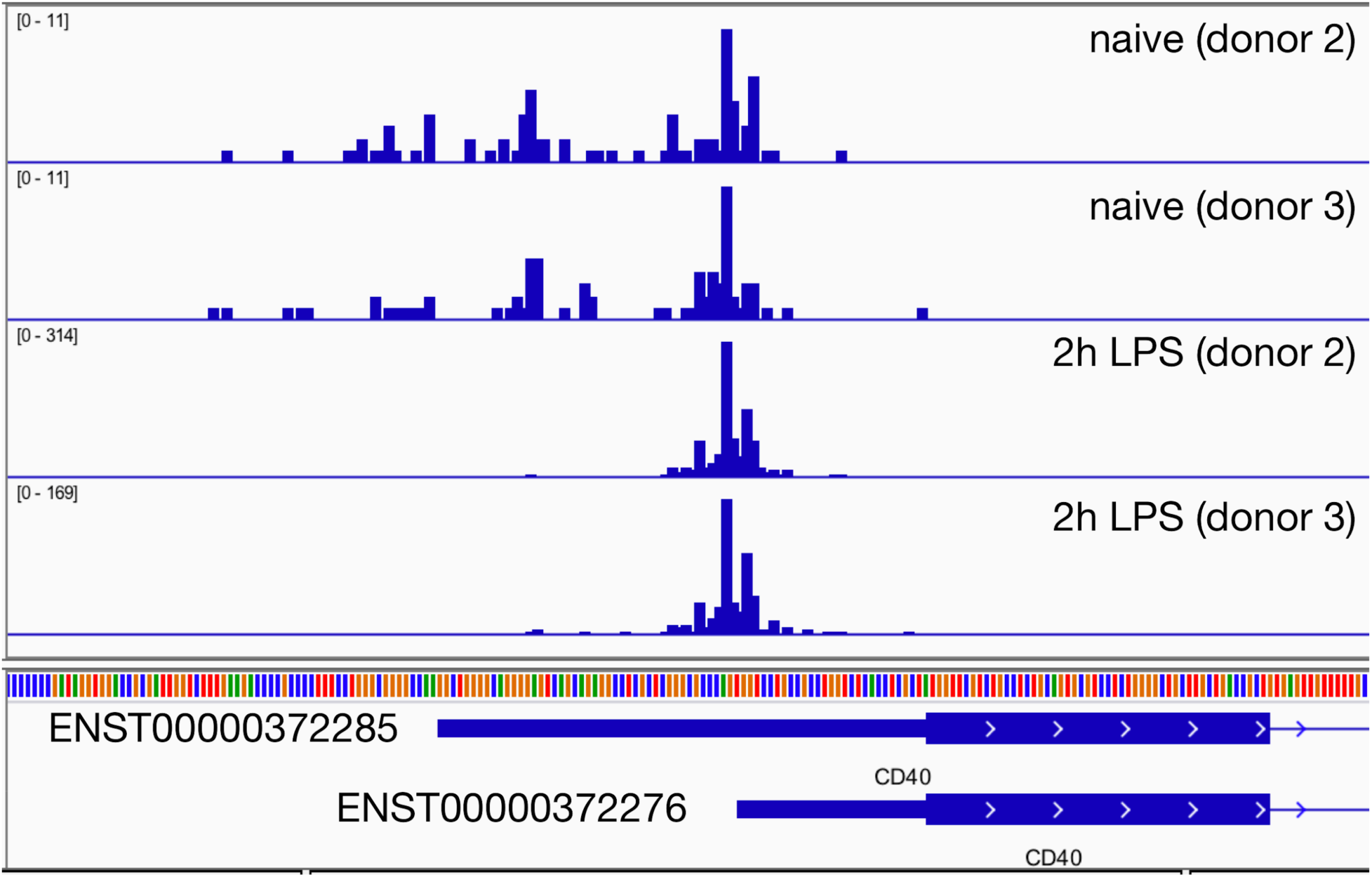
Regulation of *CD40* promoter usage in response to 2h lipopolysaccharide (LPS) in primary macrophages. The genome browser screenshots show read coverage of the Cap Analysis of Gene Expression (CAGE) data from two replicates of monocyte-derived macrophages before and after stimulation with LPS. The data was generated by the FANTOM5 consortium [37]. The *CD40* promoter containing the short 5′ UTR is strongly upregulated after 2h LPS stimulation.

